# Plant-derived coumarins shape the composition of an *Arabidopsis* synthetic root microbiome

**DOI:** 10.1101/485581

**Authors:** Mathias J.E.E.E. Voges, Yang Bai, Paul Schulze-Lefert, Elizabeth S. Sattely

## Abstract

Which factors dictate the composition of the root microbiome and its role in plant fitness is a long-standing question. Recent work has highlighted a major contribution of the soil inoculum in determining the composition of the root microbiome. However, plants are known to conditionally exude a diverse array of unique secondary metabolites, largely varying between species and environmental conditions. Here, we explore the role of specialized metabolites in dictating which bacteria reside in the rhizosphere. We employed a reduced synthetic community (SynCom) of *Arabidopsis thaliana* root-isolated bacteria to detect community shifts that occur in the absence of the secreted small molecule phytoalexins, flavonoids, and coumarins. We find that lack of coumarin biosynthesis in *f6’h1* mutant plant lines causes a shift in the root microbial community specifically under iron deficiency. We demonstrate a potential role for iron-mobilizing coumarins in sculpting the *A. thaliana* root bacterial community by inhibiting the proliferation of a relatively abundant *Pseudomonas* species via a redox-mediated mechanism. This work establishes a systematic approach enabling elucidation of specific mechanisms by which plant-derived molecules mediate microbial community composition. Our findings expand on the function of conditionally-exuded specialized metabolites and lead to new avenues to effectively engineer the rhizosphere for improving crop growth in alkaline soils, which make up a third of total arable soils.

**Significance:** The root microbiome composition is largely determined by the soil inoculum, with a distinct contribution from the host. Yet, the molecular mechanisms with which the host influences its rhizobiome are only beginning to be discovered. Using a hydroponics-based synthetic community approach, we probe the impact of root-exuded specialized metabolites in shaping the root microbiome. We uncover a role for coumarins in structuring the rhizobiome, particularly by limiting the growth of a *Pseudomonas* strain, for which we propose a mechanism of action. Our findings support the exciting possibility that root-exuded coumarins form a part of the plant’s adaptive response to iron deficiency that goes beyond iron mobilization to modulate the rhizobiome, and highlights avenues towards engineering the rhizosphere for plant health.

## Introduction

Up to 25% of total photosynthetically fixed carbon is deposited by plant roots into the surrounding soil as diverse C-containing compounds, such as sugars, amino acids, organic acids and larger rhizodeposits^1–3^. These root exudates are thought to provide key nutrients and reducing equivalents for the sustenance and proliferation of soil microbes that are able to utilize them, possibly resulting in the enrichment of specific bacterial taxa at the plant root compared to the soil biome^4^. This “rhizosphere effect” has been observed in numerous plant and crop species, most notably in *Avena barbata* (oat)^5^, *Zea mays* (maize)^6^, *Medicago sativa*^7^ and the model plant *Arabidopsis thaliana*^8^. For example, studies in oat have revealed a positive correlation between the capacity of a microbe to utilize compounds exuded by the oat root, and the microbe’s relative abundance at this underground tissue^5^. Root microbial assemblages, in turn, affect the fitness of the plant host in varied ways, including by facilitating access to key nutrients^9^ and biocontrol activity^10^.

Besides excretion of primary metabolites and carbon-rich sugars, we and others have observed that plants conditionally exude a diverse range of unique secondary metabolites^11–13^. A number of these molecules have established roles in plant nutrient acquisition and abiotic stress tolerance. For example, in iron-limiting stress conditions many plants exude small redox-active molecules involved in mobilizing iron via chelation and subsequent reduction, such as the iron-mobilizing coumarins esculetin, fraxetin and sideretin in *A. thaliana*^12,14^. Less is understood about how secreted plant molecules might mediate cross kingdom interactions. However, lead examples demonstrate the potential for secreted plant metabolites to modulate the abundance and even the function of specific soil microbiota that in turn disadvantage or benefit growth and development of the host plant^15,16^: for example, certain flavonoids exuded by *Glycine max* and *Medicago sativa* initiate a cascade of molecular events in nitrogen-fixing rhizobia that result in the development of root nodules for bacterial accommodation that ultimately benefit both the plant and microbe^17,18^. In the well-studied Brassicaceae plant family, sulfur-containing phytoalexins limit the growth of pathogenic fungi and conditionally beneficial fungal endophytes, resulting in enhanced plant survival and fitness^13,19^. Despite the important contribution of secondary metabolites to host fitness in ecological and environmental extremes, such as fending off pathogens or nutrient acquisition, their impact on the structure of the commensal bacterial assemblage associated with plant roots remains poorly understood.

Establishing causal links between plant metabolites and microbial community shifts remains challenging. One technical issue experimentalists face is the difficulty in tracking bacterial strains by 16S rRNA gene surveys in diverse environments, such as soils. In addition, using genetic disruption in the plant host can have pleiotropic effects on plant metabolism, and thus microbial community composition. For example, in a seminal study employing *A. thaliana*, the hormone salicylic acid (SA) was shown to affect the root microbiome profile of plants grown in both soils or a gnotobiotic system by employing defense phytohormone mutant lines^20^. However, the use of phytohormone mutant lines could have indirect effects on the plant’s metabolism, resulting in root microbiome shifts with unexplained causes. Similarly, a recent study presented a possible role for scopoletin in root microbiome assembly^11^. This study was limited to describing differential abundances at the Operational Taxonomic Unit level (OTU, >97% similarity of 16S rRNA) and was thus unable to draw a mechanistic link between exudation of the metabolite of interest and a known root-colonizing bacterial isolate.

Here, we use a reduced synthetic community (SynCom) comprised of *A. thaliana* root-derived bacterial commensals to analyze whether *A. thaliana* root-secreted specialized metabolites modulate community composition. The SynCom was designed using strains that (1) span the range of taxonomic diversity found at the *A. thaliana* root, (2) can be individually tracked by 16S rRNA gene amplicon analysis, and (3) proliferate in nutrient rich media^21^. This synthetic community was used to colonize the roots of both wild-type and select biosynthetic mutant plants with genetic disruptions in several major branches of specialized metabolism that result in metabolites that are conditionally secreted from roots and known to contribute to plant fitness under various conditions^22^. Specifically, we tested mutants unable to produce flavonoids (*tt5*)^23^, tryptophan (*cyp79Bb2/b3*)^24^- and methionine-derived defense metabolites (*myb28*)^25^, and coumarins (*f6’h1*)^12^. To overcome issues following from the heterogeneity of soils and soil substitutes, we use a hydroponic growth setup that allows us to both monitor exudate composition and control nutrient levels. We find subtle shifts in lines disrupted in tryptophan-derived defense metabolites, and no significant effects for flavonoids or methionine-derived defense metabolites at the growth stage and conditions tested. In contrast, we find that catecholic coumarins induce significant community shifts when plants are cultivated in iron-deficient media. Among several changes in the abundance of specific community members, we find that a *Pseudomonas sp.* isolate is consistently affected by biosynthesis of catecholic coumarins. We further confirm sensitivity of this *Pseudomonas sp.* to catecholic coumarins *in vitro* and propose a molecular mechanism of action involving reactive oxygen species (ROS). Taken together, these data showcase a systematic approach to investigate the effect of root-exuded specialized metabolites on the rhizobiome, and for the first time reveal a role for conditionally-exuded coumarins in modulating the structure of a synthetic community at the *A. thaliana* root.

## Results

### A distinct root microbiome profile is observed for A. thaliana using a hydroponics-based gnotobiotic setup

A hydroponics-based method developed previously^12^ to characterize *A. thaliana* root exudates was used to probe the effects of root specialized metabolites on microbial communities. We designed a 22-member synthetic community from *A. thaliana* root endophyte isolates collected from loamy soil in Cologne^21^ (**Table S1**). Members of our synthetic community (SynCom) span a wide range of the diversity found in the *A. thaliana* root microbiota in terms of family-level taxonomy^26,27^ and were selected such that each can be uniquely identified by 16S rRNA amplicon profiling (**Fig. S1**).

We first investigated whether a root-mediated shift in community composition could be detected in our hydroponics setup using the SynCom (**Fig. 1**). We inoculated the growth media surrounding 13-day old axenically-grown seedlings to final composite OD_600_ of 0.005-0.01. After a further 7-8 days of growth a 10^1^-10^2^-fold increase in total microbial abundance in the hydroponic media was observed, as determined by colony-forming units compared to unplanted controls (**Fig. S2**). 16S rRNA profiling revealed significant differential relative abundances between SynCom members in the root compared to the starting inoculum. We observed some inherent stochasticity in root colonization by some members between experimental batches with independently cultured and assembled SynComs. Nevertheless, of the 9 strains that consistently yielded appreciable reads in all samples, we found that Proteobacteria, such as *Rhizobium sp.* Root149, *Pseudomonas sp.* Root329, and *Sinorhizobium sp.* Root1312 were consistently enriched at the root, while the Bacteroidetes and Firmicutes isolates were not (**Fig. 1b and Fig. S2**). While we could not exclude effects of the media composition on growth of these strains, our observation that Proteobacteria are among the strains most abundant at the *Arabidopsis* root is consistent with community composition analyses of plants grown in natural soils and artificial soil substrates^21,26,27^.

**Figure 1.**
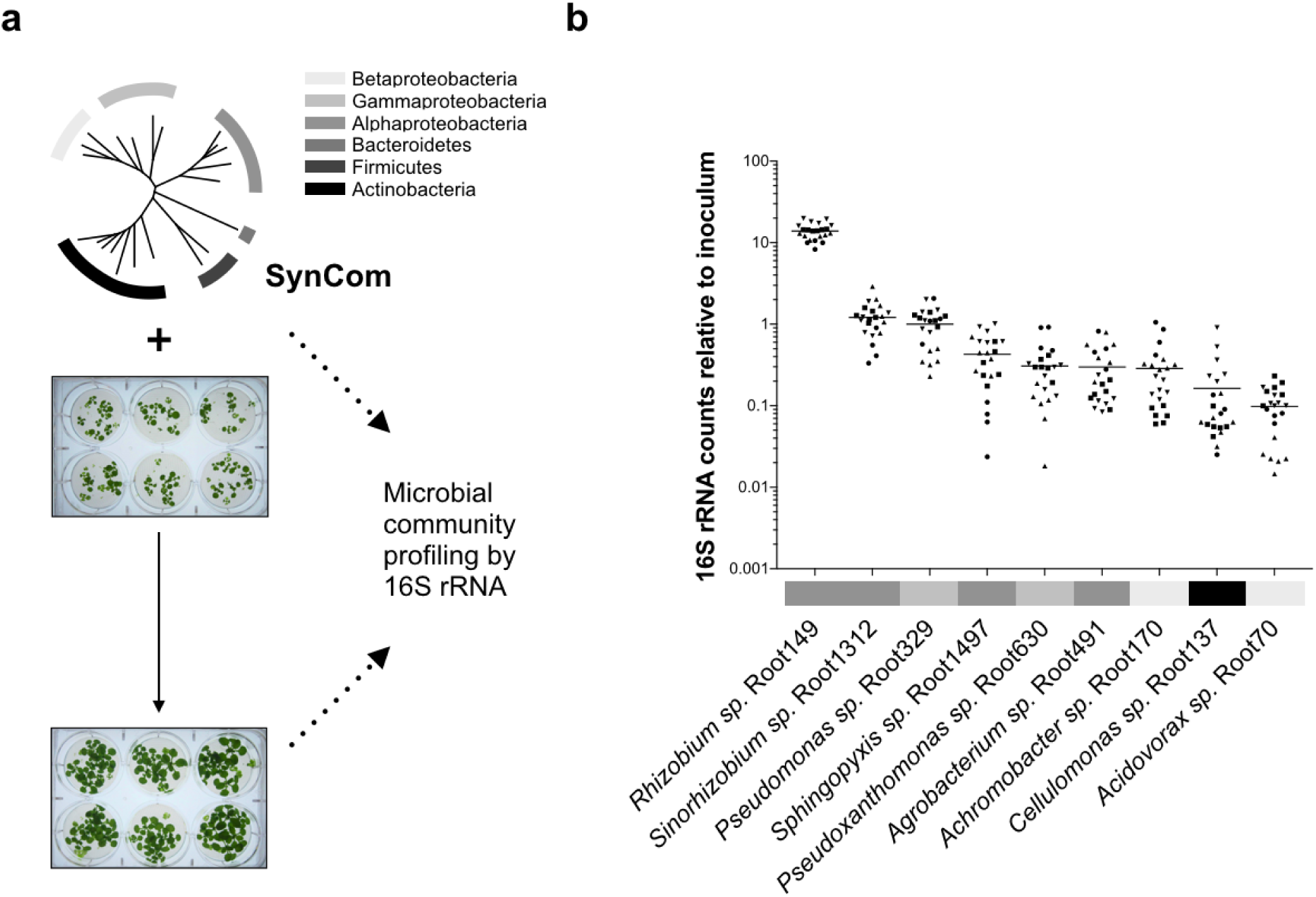
A gnotobiotic system to investigate the effects of plant metabolism on the root microbiome composition. (**a**) The media of hydroponically grown 13-day-old *A. thaliana* seedlings was inoculated with a reduced SynCom (see Fig. S1 and Table S1 for composition and properties of SynCom). The community profile at the 20-day-old root was assessed by 16S rRNA along with the starting inoculum. (**b**) Scatter plot with averages of each SynCom member’s abundance at the root relative to the starting inoculum as determined by 16S rRNA profiling. Isolates are sorted in descending order based on abundance relative to inoculum. Abundances were calculated from 16S rRNA counts (rarefied to 10^4^ reads) of four independent experiments (indicated by the symbol). Each data point represents pooled tissue harvested from two sample wells, each 16 seeds planted. Sample values >0.1 suggest enrichment of isolate at the root, as >10-fold increase in colony-forming units (CFUs) is observed in planted media at time of harvest (see Fig. S2a). Only SynCom members for which 16S rRNA reads were consistently retrievable from the inoculum and root are shown (see Fig. S2b). Bacterial phyla are indicated in accordance with the legend in (a).

### Strain relative abundances in roots of biosynthetic mutants grown in nutrient-replete growth conditions

Indole- and methionine-derived defense metabolites, flavonoids and coumarins (**Fig. 2a**) have previously been identified in *A. thaliana* root exudates and found to vary across ecotypes and species^12,16^. These studies suggest that molecules derived from these branches of specialized metabolism play a possible role in the local adaptation of the plant to the soil environment and microbial ecology. Furthermore, transcriptomics of the *A. thaliana* root colonized by commensal bacteria revealed upregulation of genes involved in the biosynthesis of these four branches of specialized metabolism^28^. To investigate the possibility that these specialized metabolites have a role in shaping the root microbiome, we determined the SynCom composition across *A. thaliana* lines that carried mutations in pathway genes required for metabolite biosynthesis in comparison to wild-type plants. We tested plant lines disrupted for: cytochrome P450 79B2 (CYP79B2) and cytochrome P450 79B3 (CYP79B3), known to initiate the indole glucosinolate and camalexin biosynthetic pathways^24^; MYB28, the positive transcriptional regulator of aliphatic glucosinolates^25^; chalcone isomerase (*tt5*), the committed step towards flavonoid biosynthesis^23^; and Feruloyl-CoA ortho-hydroxylase 1 (F6’H1), the committed step towards oxidized coumarin biosynthesis^14^ (**Fig. 2a**). We employed analysis of similarity (ANOSIM) on the Bray-Curtis dissimilarity matrix to assess significance of community composition shifts across mutant genotypes. Overall, we did not observe significant changes in the microbial composition of either *cyp79b2/b3*, *tt5, myb28* or *f6’h1* compared to wild-type in nutrient replete conditions (here defined as *p*<0.01 in ANOSIM). This finding is visually depicted in the degree of sample overlap in non-metric multidimensional scaling (NMDS) ordination plots (**Fig. S3**). To determine strain-level variation, we examined the abundance of individual isolates using Z scores. Here, the Z score was defined as: the differences between read counts for isolates from the roots of mutant lines in each sample and corresponding isolates from synchronously-grown wild-type roots, normalized to the standard deviation of read counts for wild-type root isolates (Z = *x*_i_mutant_ -μ_i_WT_)/σ_i_WT_). This analysis revealed a significant decrease in the relative abundance of *Achromobacter sp.* Root170 in *cyp79b2/b3*, and an increase of *Acidovorax sp.* Root70 in *myb28* (**Fig. 2b;** see methods section for more details on Z score calculations).

**Figure 2.**
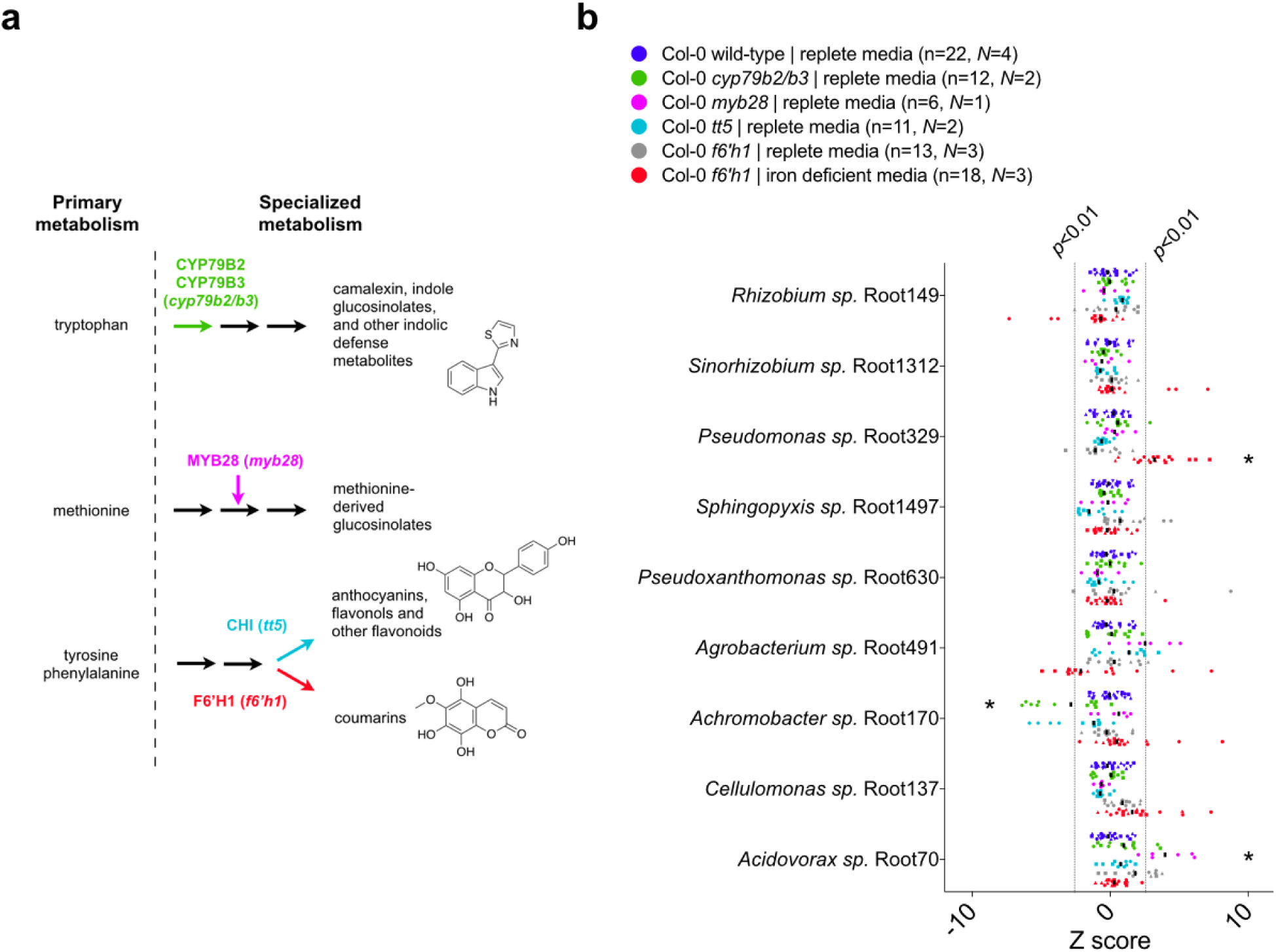
Root microbial community composition across *A. thaliana* gene disruption lines lacking major specialized metabolites. (**a**) Schematic of *A. thaliana* metabolism highlighting major pathways towards root-exuded specialized metabolites. Molecules representative of specialized metabolism are shown; these are camalexin, kaempferol and sideretin (top-down). The genes investigated are indicated by different colors with the designation used for corresponding gene-disruption lines given between brackets (see Table S2 for specific plant lines used). (**b**) Scatter plot of Z scores. Here, the Z score was defined as: the differences between read counts for isolates from the roots of mutant lines in each sample and corresponding isolates from synchronously-grown wild-type roots, normalized to the standard deviation of read counts for wild-type root isolates (Z = *x*_i_mutant_ - μ_i_WT_)/ σ_i_WT_). Asterisks indicate strains that consistently differ for that genotype. Isolates are sorted (top-down) based on abundance relative to the inoculum. Z scores corresponding to *p*<0.01 and *p*>0.01 are highlighted, and the median is indicated by vertical black bars. Z scores were calculated from 16S rRNA counts (rarefied to 10^4^ reads) of *N* independent experiments (indicated by different symbols). Each data point (n) represents pooled root tissue harvested from two sample wells, each with 16 seeds planted.

### A unique community composition shift is observed for plant lines lacking coumarin biosynthesis when grown under iron-deficiency

Oxidized coumarins downstream of scopoletin, such as esculetin, fraxetin and sideretin, are typically synthesized and exuded at appreciable levels when *A. thaliana* experiences iron deficiency to aid in the mobilization of biologically inaccessible forms of iron^12^ (**Fig. 3a**). F6H1 represents the dedicated biosynthetic step to coumarin production; mutations in this gene are unable to produce all coumarins show in **Fig. 3a**^14^. To investigate whether these oxidized coumarins affected the SynCom composition we compared microbial community profiles of *f6’h1* and wild-type in iron-deficient growth conditions (**Fig. 2b**). While there was no significant shift in community profile between wild-type and *f6’h1-1* in nutrient-replete conditions, we observed community profiles distinct from wild-type in iron-deficient conditions (ANOSIM *p*<0.01) (**Fig. S4**). Inspection at the strain level revealed a consistent increase in the relative abundance of *Pseudomonas sp.* Root329, and a minor, though not significant, decrease of *Agrobacterium sp.* Root491 in *f6’h1* lines compared to wild-type (as determined by Z score analysis; **Fig. 2b**). In line with the importance of oxidized coumarin release, we observed community profiles similar to *f6’h1* in *pdr9* (PLEIOTROPIC DRUG RESISTANCE 9) disruption mutants (**Fig. 3a-b**). The ABC transporter PDR9 has been shown to be required for the selective exudation of catecholic coumarins (esculetin, fraxetin, sideretin) under iron deficiency in *Arabidopsis* roots^29^.

**Figure 3.**
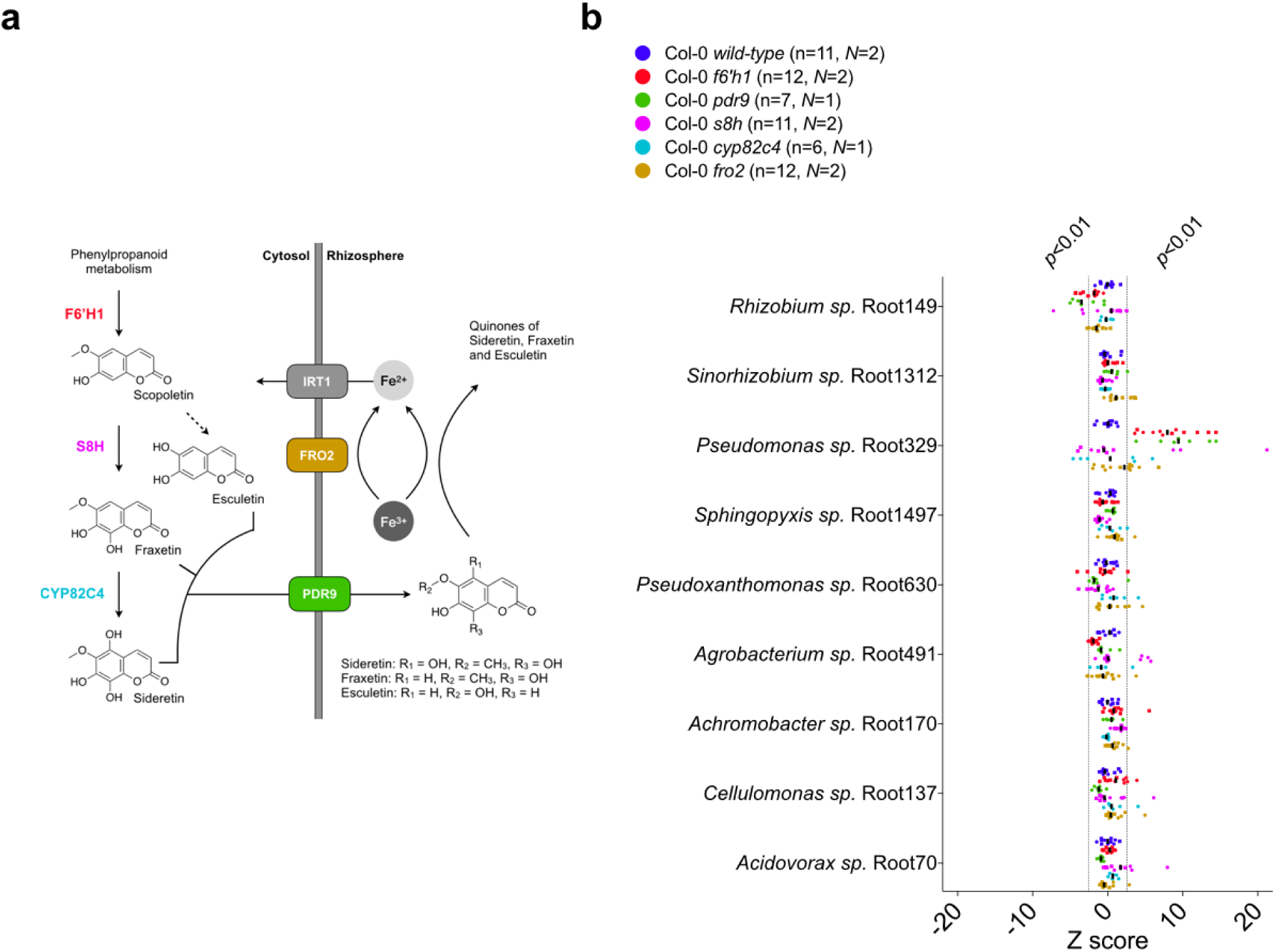
Root-exuded catecholic coumarins uniquely shift the microbial community profile at the root. (**a**) Schematic of the *A. thaliana* iron homeostasis genetic repertoire, including genes involved in oxidized coumarin biosynthesis and release. Sideretin, fraxetin and esculetin are confirmed major coumarins exuded by wilt-type roots under iron deficiency at circumneutral pH. (**b**) Scatter plot of Z scores (calculated as described in Fig. 2). Isolates are sorted (top-down) based on abundance relative to the inoculum. Z scores corresponding to *p*<0.01 and *p*>0.01 are highlighted, and the median is indicated by vertical black bars. Z scores were calculated from 16S rRNA counts (rarefied to 10^4^ reads) of *N* independent experiments (indicated by different symbols). Each data point (n) represents pooled root tissue harvested from two sample wells, each with 16 seeds planted.

To confirm that the observed community shift was due to catecholic coumarins, and not by secondary effects brought on by impaired host iron homeostasis in *f6’h1* lines, we included a *fro2* line for root community analysis. Ferric chelate reductase 2 (FRO2) is the principal ferric chelate reductase at the *A. thaliana* root^30^ (**Fig. 3a**). Mutant *fro2* lines present with severe iron deficiency symptoms in alkaline soils with low iron bioavailability. We find that community profile shifts in *f6’h1-1* lines are significantly dissimilar from wild-type plants, whereas the community profiles of *fro2* lines are not (**Fig. 3b, Fig. S5**), suggesting that the observed community profile shifts are specific to the lack of coumarin biosynthesis in *f6’h1-1*, rather than impaired host iron homeostasis.

Next, we compared the root community profiles of insertion mutants in scopoletin-8-hydroxylase (S8H) (*s8h*) and in cytochrome P450 82C4 (CYP82C4) (*cyp82c4*) with wild-type to determine whether any particular coumarin type is exclusively involved in the community shifts observed in *f6’h1* (**Fig. 3a**). The primary coumarin types exuded under iron deficiency differs for each of the mutant lines tested, with wild-type exuding primarily sideretin, fraxetin and esculetin, *cyp82c4* primarily fraxetin and esculetin, and *s8h* mainly esculetin in our hydroponic growth conditions^12^ (**Fig 3a**). We find that *s8h* and *cyp82c4* plants assemble a root community profile distinct from *f6’h1-1* or *f6’h1-2*, and more similar to wild-type, suggesting that exudation of known *A. thaliana* biosynthesized catecholic coumarins is sufficient to bring about the community profile observed in wild-type plants (**Fig. 3b, Fig. S6**).

### Lack of catecholic coumarin biosynthesis and release increases the relative abundance of Pseudomonas sp. Root329

Upon closer inspection of the relative abundances for individual bacterial strains at the root of wild-type and *f6’h1*, we found that the Gamma-Proteobacterium *Pseudomonas sp.* Root329 consistently responded positively to lines that lack biosynthesis and release of oxidized coumarins. In contrast, some root-colonizing bacteria tended towards an unchanged, or in some cases, a negative response (such as for *Agrobacterium sp.* Root491; **Fig 3b, Fig 4a**). This consistency suggested that the shift in *Pseudomonas sp.* Root329 may directly result from the absence of coumarins, and not from microbe-microbe interactions, as such interactions may present with more stochasticity. Further supporting this, we observed that increases in the relative abundance of *Pseudomonas sp.* Root329 in *f6’h1* samples were compensated for by decreases in *Rhizobium sp.* Root149, *Agrobacterium sp.* Root491 and/or *Pseudoxanthomonas sp.* Root630, depending on the replicate sample (**Fig. S7c**). In addition, *in silico* rescaling without *Pseudomonas sp.* Root329 16S rRNA reads revealed a loss of a community shift between wild-type and *f6’h1* (**Fig. S7a-b**).

**Figure 4.**
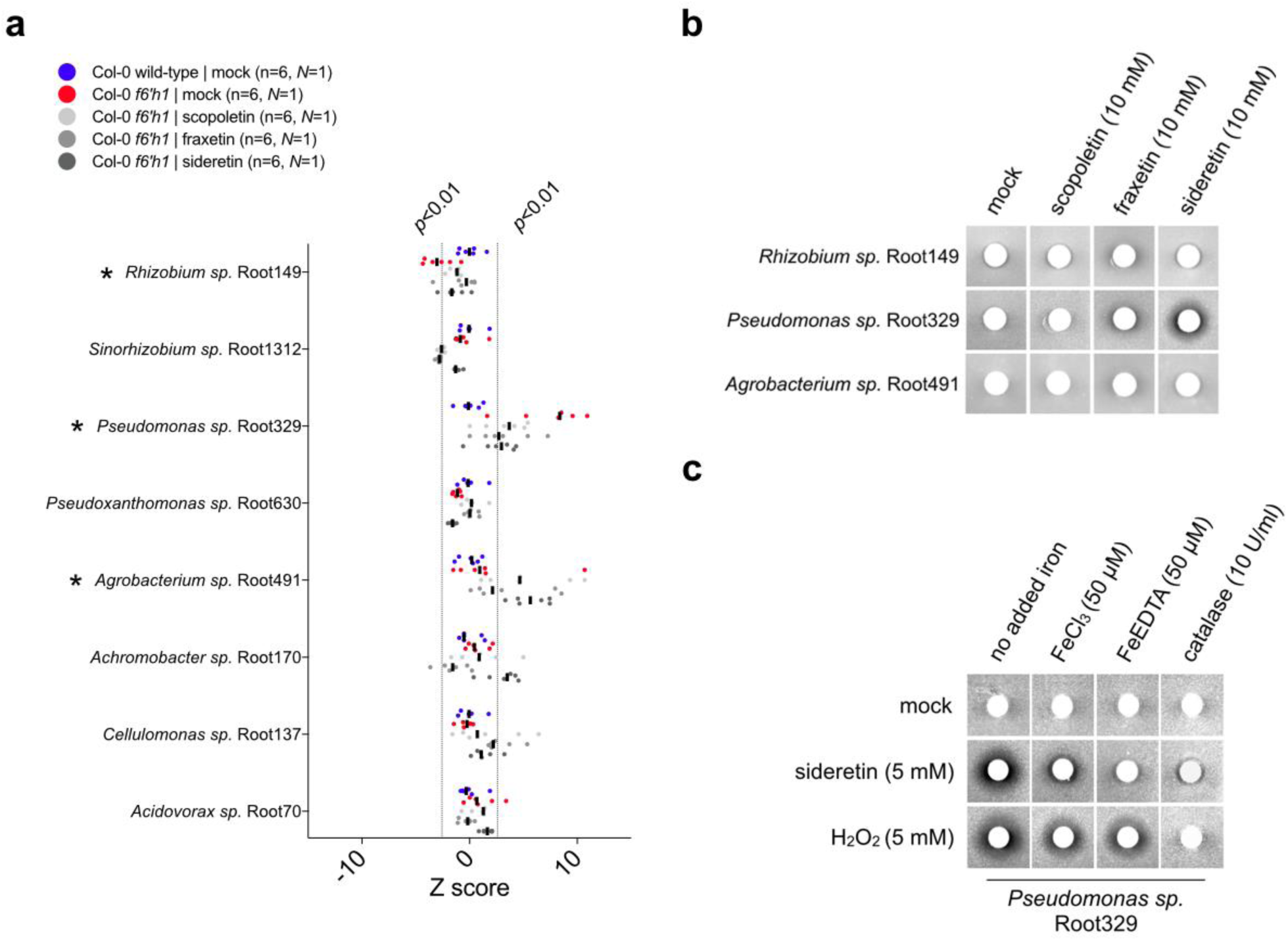
Purified coumarins partially shift the community to the composition found in wild-type plants, and specifically inhibit the growth of the differentially abundant *Pseudomonas sp.* Root329 isolate via an ROS-mediated mechanism. (**a**) Addition of 100 µM oxidized coumarins to media of *f6’h1* lines phenocopies wild-type community composition. Z scores calculated for each isolate’s RA at the mutant plant root compared to the RA at synchronously-grown wild-type roots. Z scores corresponding to *p*<0.01 and *p*>0.01 are highlighted, and the median is indicated by vertical black bars. Z scores were calculated from rarefied 16S rRNA counts. Each data point (n) represents pooled root tissue harvested from two sample wells, each with 16 seeds planted. (**b**) Assays for *in vitro* bacterial growth inhibition surrounding 5 mm filter discs (white circles) impregnated with DMSO (control), 10 mM scopoletin, fraxetin or sideretin (all in 10% DMSO). Bacteria were inoculated into nutrient-agar and impregnated filter discs were placed on top of agar. Images were taken after 48 hours growth at 30 C. The extent of the dark zones surrounding the filter discs represents the degree of bacterial growth inhibition. The experiment was repeated three times with similar results. (**c**) Bacterial growth surrounding filter discs impregnated with DMSO (control), sideretin or hydrogen peroxide at 5 mM (all in 10% DMSO). Conditionality of sideretin-mediated growth inhibition was assessed by addition of insoluble iron (FeCl_3_), soluble iron (Fe-EDTA) or catalase to nutrient-agar plates prior to inoculation.

### Coumarin complementation of f6’h1 phenocopies the wild-type SynCom profile

Next, we assessed the microbial community profiles of *f6’h1* lines grown in the presence of 100 µM scopoletin, fraxetin or sideretin under iron-deficient conditions (**Fig 4a**). The addition of either oxidized coumarin was sufficient to phenocopy wild-type SynCom profiles. Most notably, shifts in the relative abundance of *Rhizobium sp.* Root149 and *Pseudomonas sp.* Root329 were lost, with the exception of *Agrobacterium sp.* Root491, which responded positively to the addition of oxidized coumarins both in wild-type and *f6’h1* plants. Appreciable levels of sideretin was observed in the media of *f6’h1* plants that were chemically complemented with scopoletin, consistent with previous studies^14^, suggesting that the community shifts observed may not be caused directly by scopoletin (as scopoletin could be converted to fraxetin and/or sideretin in *f6’h1* plants).

### Growth of Pseudomonas sp. Root329 is inhibited by iron-mobilizing coumarins in vitro

To determine if F6’H1-dependent coumarins directly impact the growth of bacterial isolates that comprise the SynCom, we analyzed the sensitivity of the major coumarin-responsive root-proliferating SynCom members to purified samples of scopoletin, fraxetin and sideretin *in vitro*. Notably, using a halo inhibition assay we observed severe growth inhibition of *Pseudomonas sp.* Root329 by the predominant wild-type coumarin sideretin, and moderate inhibition by fraxetin (**Fig 4b**). The differentially abundant strains *Rhizobium sp.* Root149, and *Agrobacterium sp.* Root491 were not comparably affected by either coumarin; however, in some instances, a faint ring of apparent growth inhibition was observed surrounding fraxetin-impregnated discs.

We further dissected the mechanism of action for catecholic coumarins on *Pseudomonas sp.* Root329. We hypothesized that some oxidized coumarins’ molecular structures – harboring a highly reduced catechol moiety – allowed for the reduction of oxygen to reactive oxygen species (ROS), which are potent antimicrobials. In support of a model for coumarin-mediated antimicrobial activity via the ROS hydrogen peroxide, we found that the inhibitory activity of sideretin on *Pseudomonas sp.* Root329 could be abolished with addition of catalase (**Fig 4c**). Notably, the antimicrobial activity of sideretin could further be reduced by the addition of soluble iron (**Fig 4c**). These findings suggest peroxide generation by catecholic coumarins is promoted when iron is scarce – the typical environment in which iron-mobilizing coumarins are biosynthesized and secreted by the *A. thaliana* root. In addition, we observed hydrogen peroxide generation by the catecholic coumarins sideretin, fraxetin and esculetin (but not for scopoletin or isofraxidin, which lack catechol functionality) in iron-deficient MS media over a 24-hour period (**Fig. S9**). Sideretin was the most potent hydrogen peroxide generator at shorter time scales, which may explain its antimicrobial effect on *Pseudomonas* Root329 in the LB-agar assay performed over a 24-hour timespan. Further, in support of a ROS-mediated mechanism of inhibition, we challenged the most abundant SynCom members at the root with hydrogen peroxide and found that *Pseudomonas sp.* Root329 was most affected (based on CFU counts) (**Fig. S10**).

## Discussion

Studies into the host factors that shape the *A. thaliana* rhizosphere community structure have been few, with pioneering work showing that salicylic acid is an important player modulating root community composition^20^. In contrast to phytohormones, a recent study has correlated microbial growth at the root of oat to the microbe’s ability to utilize compounds found to be exuded^5^. In general, it was found that utilization of aromatic amino acids is highly correlated with growth at the root. These examples provided impetus for investigating the role of the broader range of diverse metabolites exuded from plant roots on the composition and function of the root microbiome.

Here, we employed a synthetic community composed of *A. thaliana* natural root isolates to investigate the importance of specialized metabolites on microbial community composition at the *A. thaliana* root. Of the mutant lines investigated, we find significant changes in relative microbial abundances for coumarin biosynthesis mutants, specifically under iron-limiting conditions. Notably, we find an average 3-fold increased abundance of *Pseudomonas sp.* Root329 for plant lines devoid of coumarin biosynthesis or exudation and provide evidence that this change in relative abundance could be due to this strain’s sensitivity to the major catecholic coumarins produced by iron-deficient wild-type plants. As a reference, the changes in relative abundances observed in this study are on par with those observed in previous studies utilizing synthetic communities. For example, an increase in *Terracoccus sp*. RA of about two fold, and a three-fold decrease in *Mitsuria sp.* RA upon addition of salicylic acid (SA) to the roots of *Arabidopsis* has been shown to correspond to these strain’s growth-response to SA *in vitro*^20^.

Our study provides direct evidence that specialized metabolites can cause reproducible changes in community composition of plant roots. While these interactions are likely much more complex in soil grown plants, the hydroponic system using a defined bacterial community provides clear evidence that relatively small changes in the secretion of specialized metabolites could feasibly impact community structure – even in the soil. For example, in light of our data, one possible explanation for the observed variation of communities in the roots of *f6’h1* plants grown in natural soil^11^ compared to those of wild-type can be explained by the major catecholic coumarins produced under iron limitation: esculetin, fraxetin and sideretin.

Based on our findings, we propose that the catechol moiety of iron-mobilizing coumarins promotes redox reactions that can both mobilize iron via reduction, and/or generate antimicrobial reactive oxygen species, such as hydrogen peroxide, with detrimental effects on microbial proliferation. Similar antimicrobial mechanisms of action have been described for diverse phenazines produced by some Pseudomonads – these redox-active small molecule’s functionality (either generation of ROS or oxidation by metal ions) is dictated by the surrounding iron and oxygen levels^31^. Furthermore, ROS have been implicated in the mechanism of action for a number of antimicrobials, suggesting that the mechanisms underlying tolerance to catecholic coumarins may be similar to those of known antimicrobials^32^.

Bacteria harbor sophisticated mechanisms for iron mobilization and sequestration that, in turn, may have multifaceted effects on host fitness^33,34^. It is possible that root bacterial community shifts brought on by iron-mobilizing coumarins could bestow an adaptation to the host plant experiencing iron-deficiency, for example by specifically fending off strong microbial competitors for iron. In addition, root-associated bacteria may have evolved mechanisms of coumarin sensing that prove adaptive to the host plant experiencing iron-deficiency symptoms. We explored this possibility using an agar plate-based assay to assess seedling growth upon inoculation with SynCom members under iron-deficiency. However, while we observed seedling growth promotion by the complete SynCom for wild-type plants (**Fig. S11**), these effects were also conferred by the majority of individual SynCom members in mono-association experiments (with the exception of the Firmicutes, which did not proliferate at the root; **Fig. S12**). Furthermore, we observed no significant difference in growth promotion of wild-type and *f6’h1* seedlings between *Pseudomonas sp.* Root329, *Rhizobium sp.* Root149 (a highly-abundant isolate that did not display significant RA shifts in *f6’h1*), or plants inoculated with equal densities of both strains (**Fig. S13**). While this assay does not reveal an obvious direct benefit, it is still possible that the host-induced community shifts could confer an adaptive advantage using assays for plant growth and development in less artificial conditions. Furthermore, future work should investigate the transcriptional response of purified microbial isolates to single coumarin types to identify the metabolic and/or signaling pathways are responsive to coumarins, which may offer insights into the molecular mechanisms of action underpinning seedling growth promotion by root isolates.

We demonstrate here the promise of employing genetic disruption mutants in biosynthetic pathways and reduced synthetic communities to identify the specific molecular players that sculpt the rhizosphere microbiome. In particular, the ability to track individual members of the SynCom facilitates subsequent investigation of the underpinning molecular mechanism *in vitro*^35^. A deeper understanding of the molecular mechanisms required for bacterial colonization of the root and rhizosphere will facilitate efforts towards engineering host-beneficial microbial consortia. Our finding that proliferation of specific microbial strains is affected by iron-mobilizing coumarins can be used to effectively engineer the rhizosphere for improving crop growth in alkaline soils, which make up a third of total arable soils^36^.

## Supporting information

## Acknowledgements

We thank K. Huang, J. Rajniak, R. Garrido-Oter, M. Hashimoto, and H. Inoue for valuable discussions. We thank S. Long, S. Sirk, A. Denisin, A. Klein, C. Liou, R. Nett and F. Hol for valuable feedback on the manuscript. We thank X. Ji for technical assistance with Illumina sequencing at the Stanford Functional Genomics Facility. We thank H. Inoue for help developing the seedling growth promotion assays. M.J.E.E.E.V. acknowledges support from the Stanford Bio-X Graduate Fellowship. This work was supported by an HHMI and Simons Foundation Grant 55108565 and NIH DP2 Grant AT008321 (to E.S.S.).

---

**Table S1.** Taxonomy of bacterial strains that compose the synthetic community used in this study. All strains were isolated from *A. thaliana* roots grown in Cologne wild-type soil^21^.

| Strain ID | Phylum | Class | Family | Genus |
| --- | --- | --- | --- | --- |
| Root137 | Actinobacteria | Actinobacteria | Cellulomonadaceae | *Cellulomonas* |
| Root166 | Actinobacteria | Actinobacteria | Microbacteriaceae | *Microbacterium* |
| Root332 | Actinobacteria | Actinobacteria | Microbacteriaceae | *Yonghaparkia* |
| Root563 | Actinobacteria | Actinobacteria | Intrasporangiaceae | *Janibacter* |
| Root614 | Actinobacteria | Actinobacteria | Nocardiodaceae | *Nocardioides* |
| Root1310 | Actinobacteria | Actinobacteria | Streptomycetaceae | *Streptomyces* |
| Root1464 | Actinobacteria | Actinobacteria | Microbacteriaceae | *Agromyces* |
| Root186 | Bacteroidetes | Flavobacteriia | Flavobacteriaceae | *Flavobacterium* |
| Root11 | Firmicutes | Bacilli | Bacillaceae | *Bacillus* |
| Root444D2 | Firmicutes | Bacilli | Paenibacillaceae | *Paenibacillus* |
| Root483D1 | Proteobacteria | Alphaproteobacteria | Methylobacteriaceae | *Methylobacterium* |
| Root149 | Proteobacteria | Alphaproteobacteria | Rhizobiaceae | *Rhizobium* |
| Root491 | Proteobacteria | Alphaproteobacteria | Rhizobiaceae | *Agrobacterium* |
| Root1312 | Proteobacteria | Alphaproteobacteria | Rhizobiaceae | *Sinorhizobium* |
| Root1497 | Proteobacteria | Alphaproteobacteria | Sphingomonadaceae | *Sphingopyxis* |
| Root70 | Proteobacteria | Betaproteobacteria | Comamonadaceae | *Acidovorax* |
| Root170 | Proteobacteria | Betaproteobacteria | Alcaligenaceae | *Achromobacter* |
| Root189 | Proteobacteria | Betaproteobacteria | Oxalobacteraceae | *Herbaspirillum* |
| Root329 | Proteobacteria | Gammaproteobacteria | Pseudomonadaceae | *Pseudomonas* |
| Root561 | Proteobacteria | Gammaproteobacteria | Xanthomonadaceae | *Rhodanobacter* |
| Root630 | Proteobacteria | Gammaproteobacteria | Xanthomonadaceae | *Pseudoxanthomonas* |
| Root1280 | Proteobacteria | Gammaproteobacteria | Moraxellaceae | *Acinetobacter* |

**Table S2.** Homozygous T-DNA insertion plant lines used in this study. All plants were of the Columbia-0 ecotype background. SALK lines were obtained from the Arabidopsis Biological Resource Center (ABRC) and SM lines from the Nottingham Arabidopsis Stock Centre (NASC). Zygosity was confirmed using primer sequences from the SALK T-DNA primer tool.

| Mutated gene | Phenotype affected | Line | Reference |
| --- | --- | --- | --- |
| – | (wild-type) | – | F. Ausubel (gift) |
| *cyp79b2/cyp79b3*<br>(At4g39950/At2g22330) | indole phytoalexin biosynthesis | – | Zhao et al. (2002) |
| *f6'h1-1* (At3g13610) | all coumarin biosynthesis | SALK_132418C | – |
| *f6'h1-2* (At3g13610) | all coumarin biosynthesis | SALK_050137C | – |
| *myb28* (At5g61420) | aliphatic glucosinolate biosynthesis | SALK_136312C | – |
| *tt5* (At3g55120) | flavonoid biosynthesis | SALK_034145 | – |
| *fro2* (At1g01580) | principal ferric chelate reductase | CS309724 | – |
| *s8h-1* (At3g12900) | fraxetin and sideretin biosynthesis | SM_3_27151 | – |
| *s8h-2* (At3g12900) | fraxetin and sideretin biosynthesis | SM_3_23443 | – |
| *cyp82c4-1* (At4g31940) | sideretin biosynthesis | SALK_001585 | – |
| *pdr9-2* (At3g53280) | catecholic coumarins root transporter | SALK_050885 | – |

**Figure S1.**
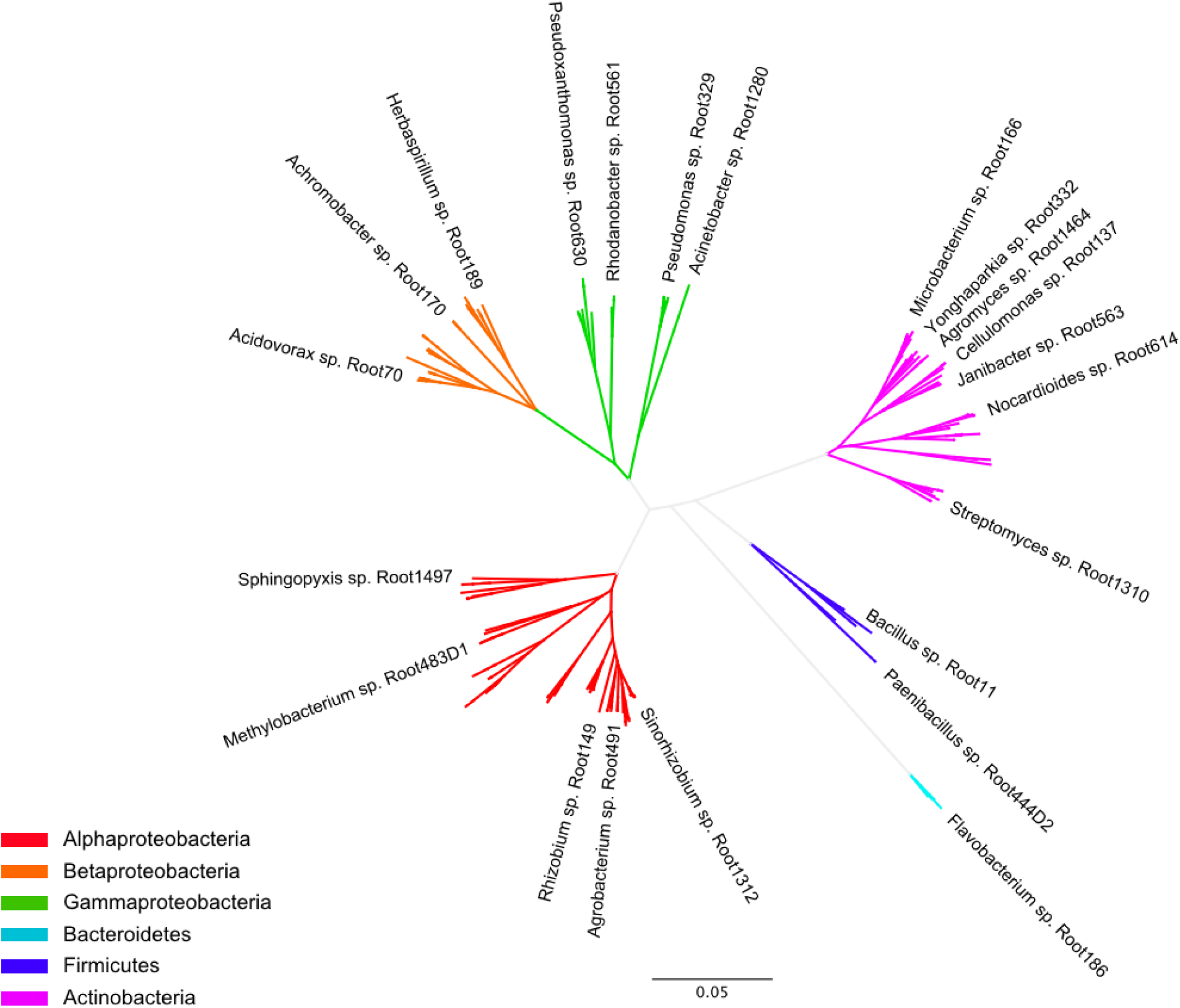
16S rRNA phylogenetic tree of bacterial strains that compose the minimal synthetic community used in this study. Genome-sequenced *A. thaliana* root isolates obtained in the study of Bai et al. (2015)^21^ are included for context (unlabeled branches). Bacterial phyla are indicated by color.

**Figure S2.**
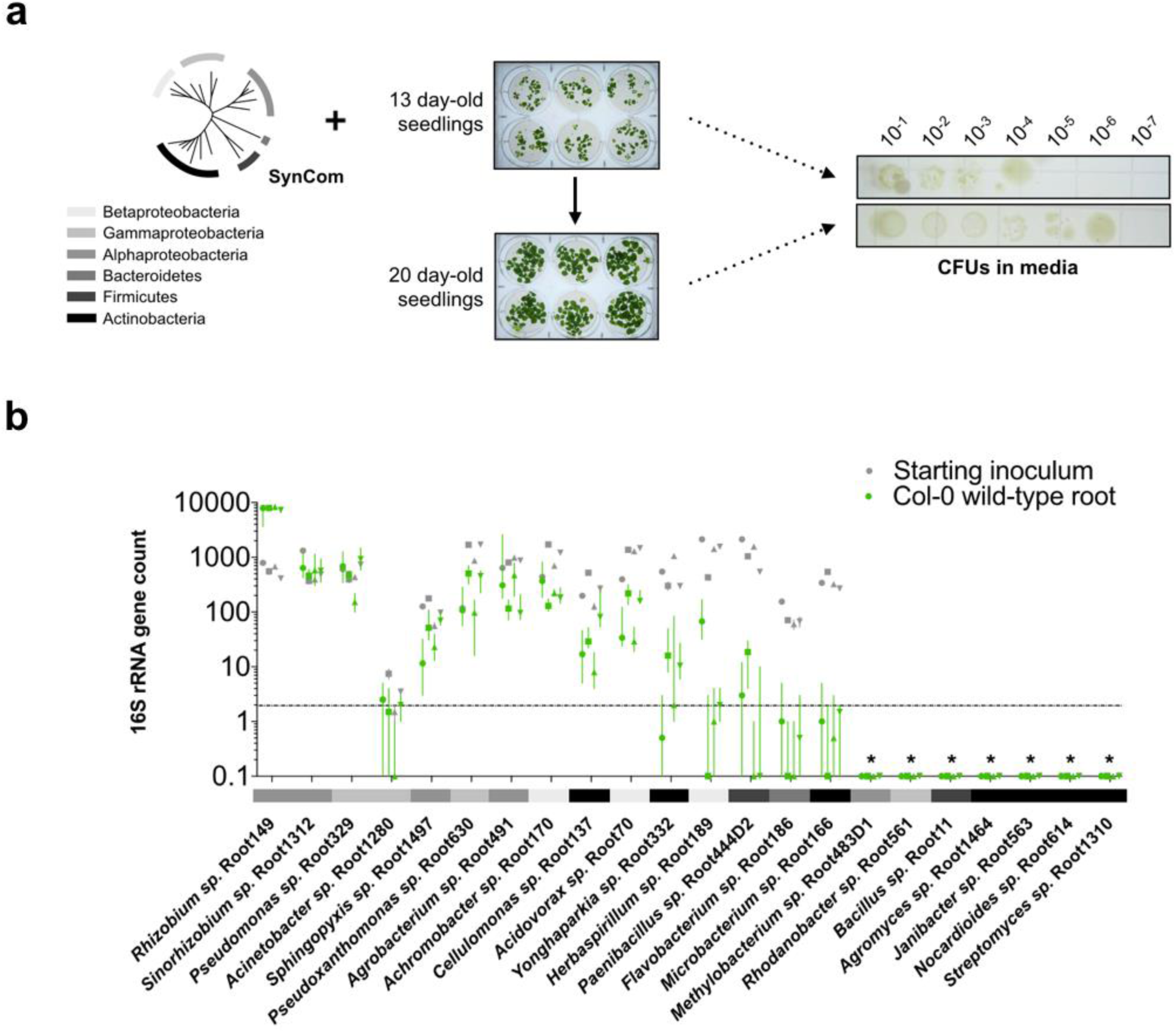
Response of SynCom members at the iron-deficient *A. thaliana* root. (**a**) Schematic as in Fig. 1a with observed CFUs in the media of 20-day old seedlings incubated with the SynCom for 7 days. Overall, a 10-fold increase in absolute abundance of the total SynCom is observed in planted media compared to the inoculum (used to inoculate 13-day-old seedlings). (**b**) Rarefied 16S rRNA counts at the wild-type root (green) and starting inoculum (grey) for the four independent experiments shown in Fig. 1b (each independent experiment represented using a different symbol). Each data point represents 2 sample wells, each with 16 planted seeds. The dashed horizontal line denotes a threshold of 2, below which data for that isolate was not further considered. Asterisks denote SynCom strains for which 16S rRNA amplicons were not retrieved, likely due to insufficient lysis of cells using standardized homogenization procedures. Phyla colors are the same as in (a).

**Figure S3.**
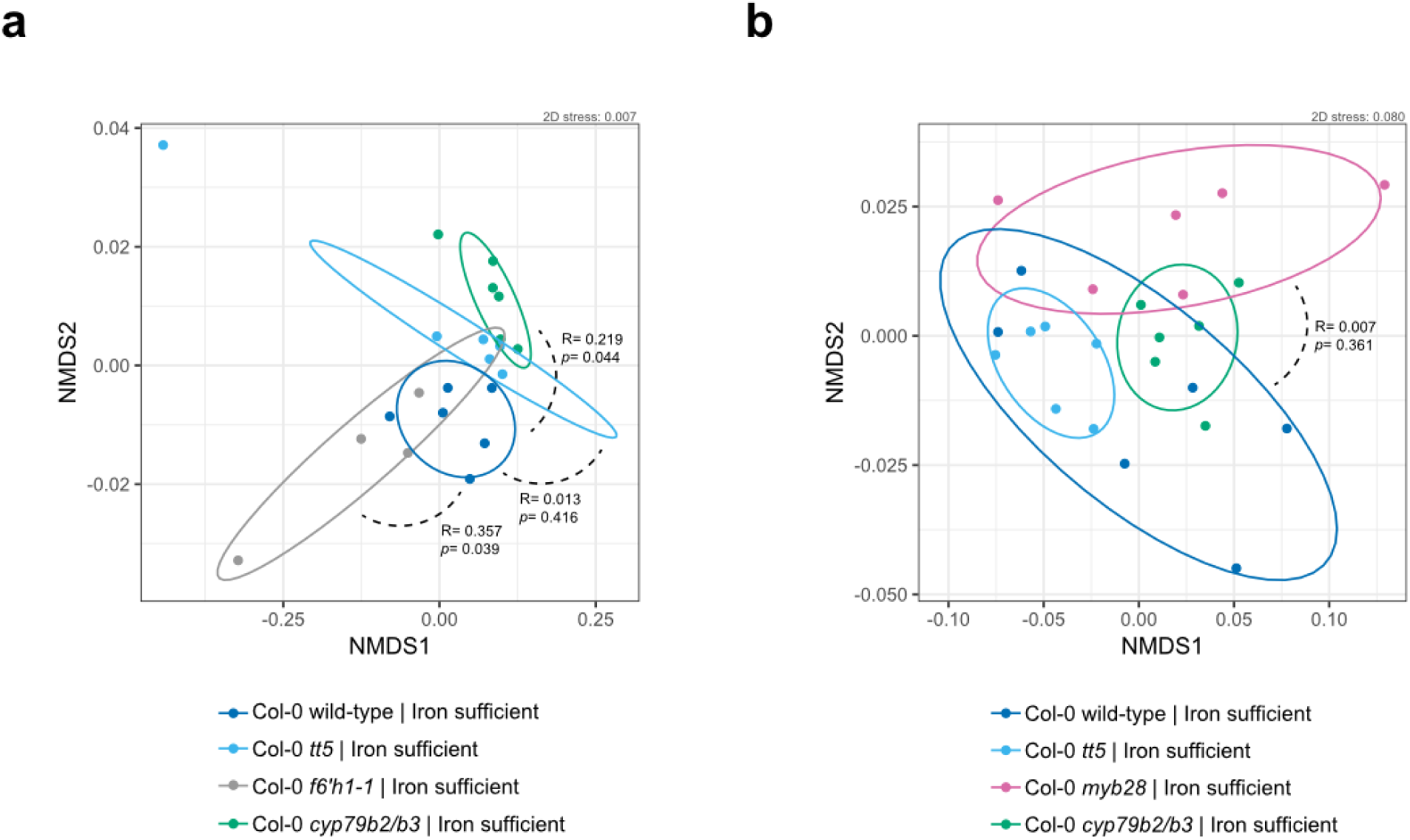
Shifts in root community profiles across *A. thaliana* gene-disruption mutants involved in major pathways of specialized metabolism. (**a**) Non-metric multidimensional scaling (NMDS) on Bray-Curtis dissimilarity matrices of microbial community profiles at 20-day-old roots of Col-0 wild-type, *tt5*, *f6’h1* and *cyp79b2/b3* (n= 22 microbial community profiles). Ellipses (75% CI) roughly depict the overlap between microbial communities of genotypes: where there is overlap there is no significant community profile shift. Degree of dissimilarity (R, low = 0, high = 1) and significance (*p*) are obtained using analysis of similarity (ANOSIM) of gene-disruption line community profiles versus wild-type community profiles. Each data point was obtained by sampling from pooled roots grown in two wells (32 planted seeds total). n= 6 for wild-type and *cyp79b2/b3*, n= 5 for *tt5* and n= 4 for *f6’h1-1*. (**b**) An independent experiment as in (a), with Col-0 wild-type (n= 6), *tt5* (n= 6), *myb28*(n= 6), and *cyp79b2/b3* (n=6). Panels (a) and (b) show data from the same experiments as in Fig 2b, each panel represents an independently-performed experiment.

**Figure S4.**
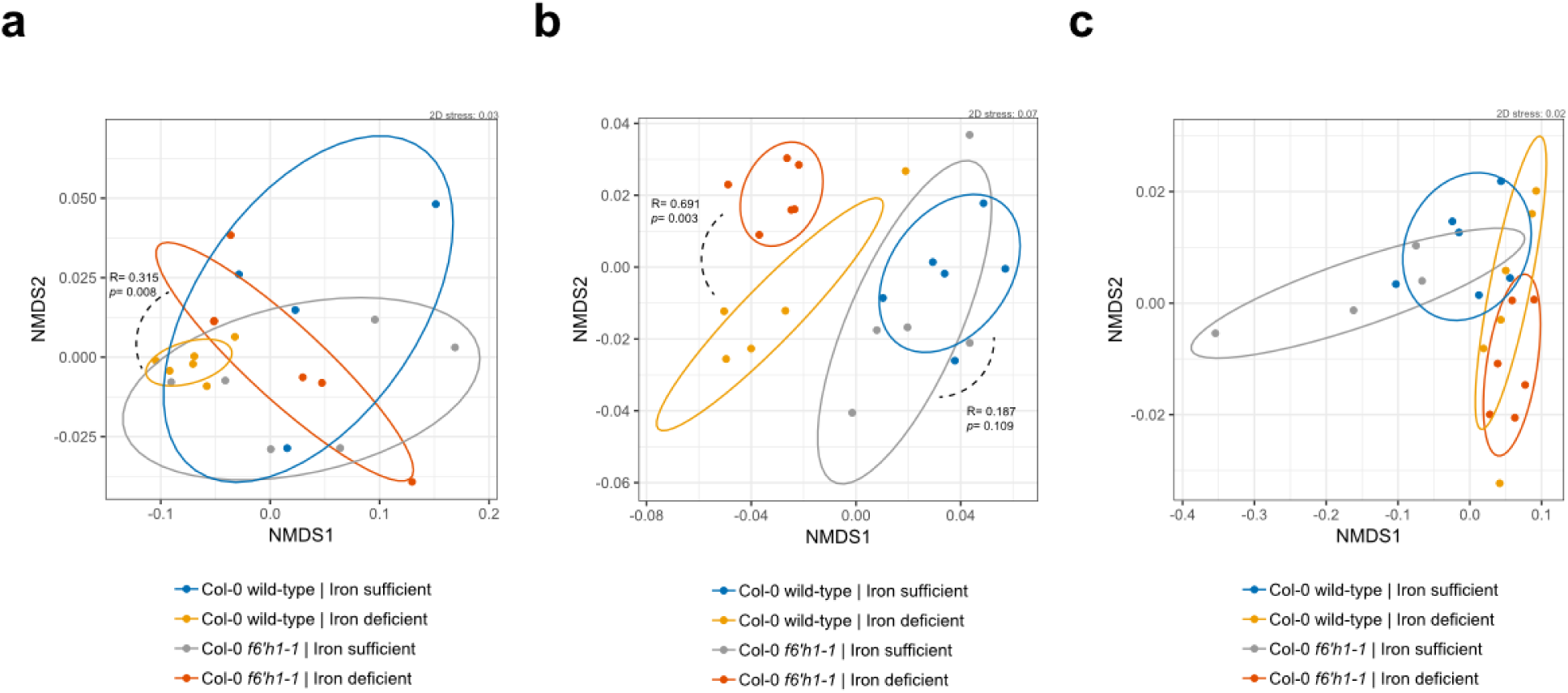
Shifts in root community profiles between *A. thaliana* Col-0 wild-type and *f6’h1-1*. (**a**) Non-metric multidimensional scaling (NMDS) on Bray-Curtis dissimilarity matrices of microbial community profiles at the wild-type and *f6h1-1* roots in nutrient replete and iron-deficient media (20-day-old seedlings). In general, there is significantly more dissimilarity between root community profiles of wild-type and *f6’h1* lines grown in iron-deficient media than those grown in replete media. Wild-type (n= 4) and *f6h1-1* (n=6) in replete media, wild-type (n= 6) and *f6h1-1* (n=6) in iron-deficient media. All other descriptors are the same as in Figure S3. (**b**) An independent experiment as in (a), with Col-0 wild-type (n= 6) and *f6’h1-1* (n=5) in replete media, and Col-0 wild-type (n= 5) and *f6’h1-1* (n=6) in iron-deficient media. (**c**) An independent experiment as in (a), with Col-0 wild-type (n= 6) and *f6’h1-1* (n=4) in replete media, and Col-0 wild-type (n= 6) and *f6’h1-1* (n=6) in iron-deficient media. Panels (a), (b) and (c) show data from the same experiments as in Fig 2b, each panel represents an independently-performed experiment.

**Figure S5.**
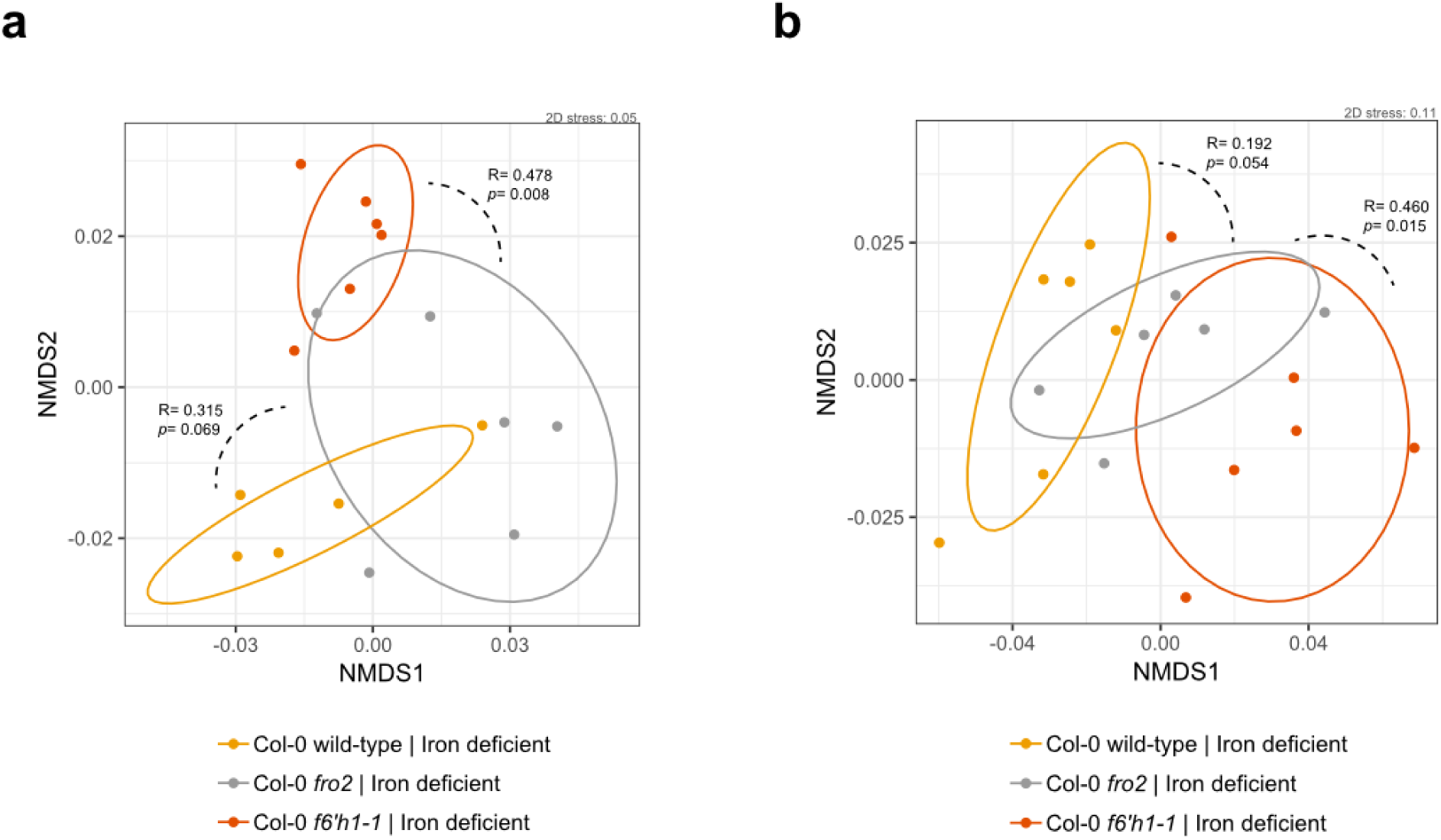
Shifts in root community profiles between *A. thaliana* Col-0 wild-type and *fro2*. Changes in root community composition between 20-day-old seedlings of *A. thaliana* Col-0 wild-type, *f6’h1-1* and *fro2* (incubated with the SynCom for 7 days) (**a-b**) Non-metric multidimensional scaling (NMDS) on Bray-Curtis dissimilarity matrices of microbial community profiles at the Col-0 wild-type, *f6h1*-1 and *fro2* roots grown in iron-deficient media (n= 17, and n=18 microbial community profiles; two independent experiments). A larger degree of dissimilarity is observed between community profiles of *f6’h1* and *fro2*, than between wild-type and *fro2,* suggesting that an impaired iron homeostasis is not the main driver of community profile shifts observed for *f6’h1*. All other descriptors are the same as in Figure S3

**Figure S6.**
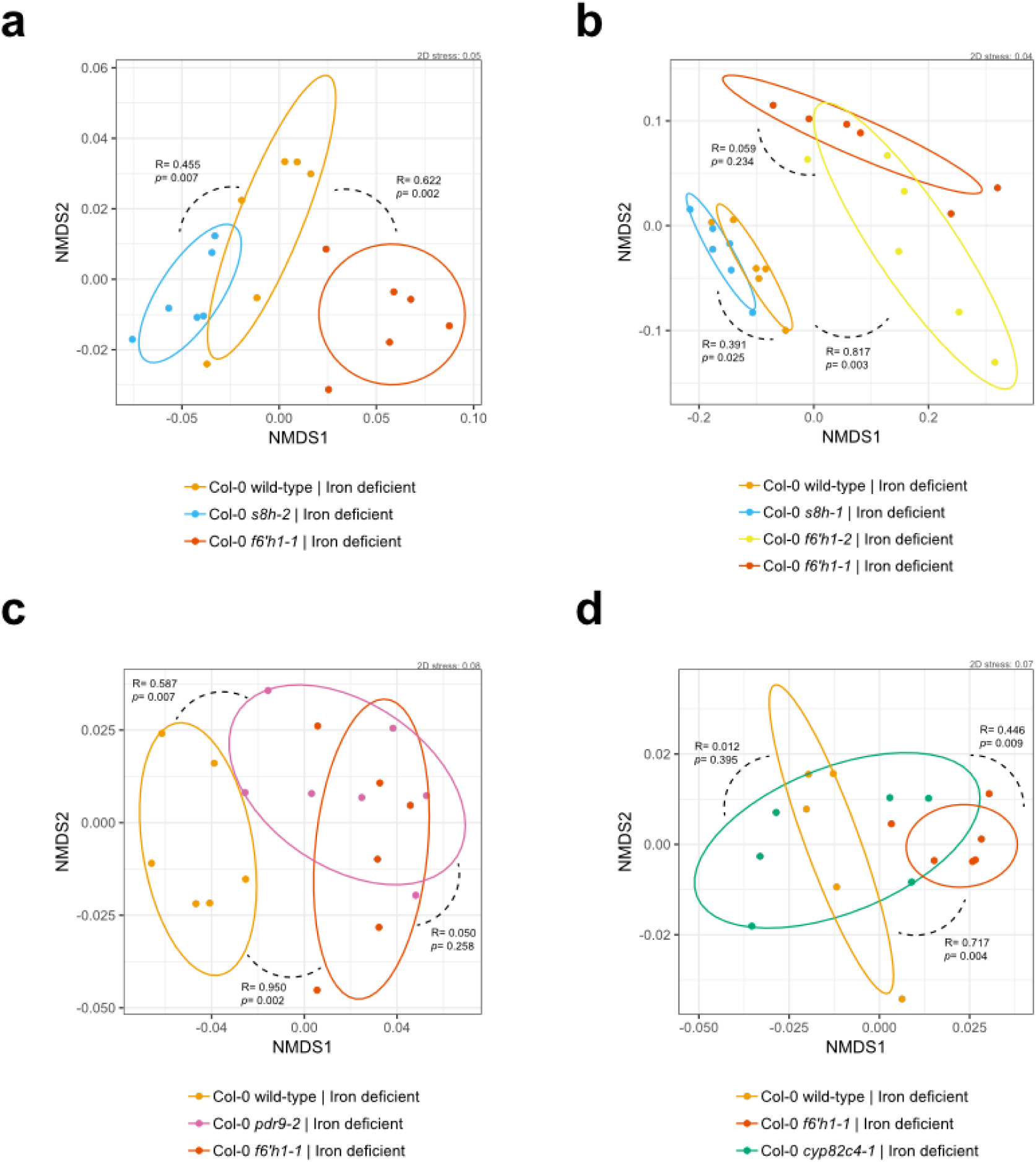
Shifts in root community profiles across coumarin biosynthesis and release mutant lines. (**a-b**) Non-metric multidimensional scaling (NMDS) on Bray-Curtis dissimilarity matrices of microbial community profiles at the wild-type, *f6’h1-1, f6’h1-2, s8h-1* and *s8h-2* roots in iron-deficient media (of 20-day-old seedlings, incubated with the SynCom for 7 days). (**c**) Same as in with wild-type, *f6h1-1* and *pdr9-2* in iron-deficient media. (**d**) Same as in (a) with wild-type, *f6h1-1* and *cyp82c4-1* in iron-deficient media. A significant degree of dissimilarity of microbial community profiles is observed between *f6’h1* and *pdr9-2* on the one hand and *s8h* and *cyp82c4-1* on the other. All other descriptors are the same as in Figure S3

**Figure S7.**
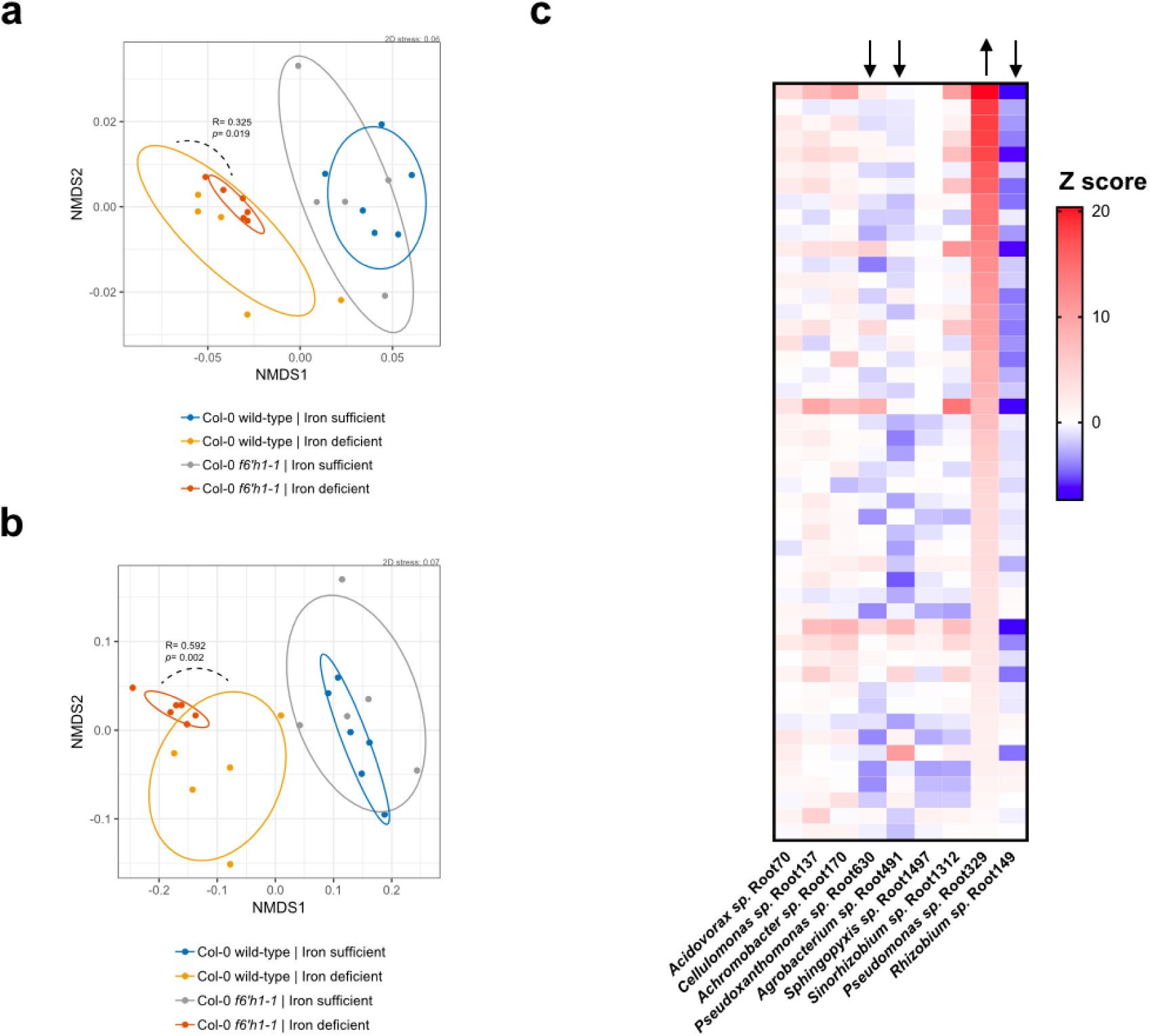
*In silico* assessment of *Pseudomonas sp.* Root329 as the main isolate responsive to oxidized coumarins. (**a**) Non-metric multidimensional scaling (NMDS) on Bray-Curtis dissimilarity matrices of microbial community profiles at wild-type and *f6h1-1* in iron-sufficient and iron-deficient roots (same experiment as in Figure S4b) with 16S rRNA reads corresponding to *Pseudomonas sp.* Root329 removed *in silico*. (**b**) Same as in Figure S4b, here rescaled without *Rhizobium sp.* Root149 RAs. By rescaling without *Pseudomonas sp.* Root329 the degree of dissimilarity between wild-type and *f6’h1-1* decreases, however removing *Rhizobium sp.* Root149 reads (largest contributor) retains the original dissimilarity. (**c**) Heatmap of each strain’s Z score (compared to wild-type) for *f6’h1* roots grown in iron-deficient media. On average, Z-scores for *Pseudomonas sp.* Root329 increase in *f6’h1* roots, without a consistent compensatory decrease for any specific strain (either *Pseudoxanthomonas sp.* Root630, *Agrobacterium sp.* Root491 and/or *Rhizobium sp.* Root149 decrease when *Pseudomonas sp.* Root329 increases; depicted by arrows). This suggest there is a stronger correlation between lack of coumarin biosynthesis and increased RA for Root329, than for (compensatory) decreased RA in any other strain. So, *Pseudomonas sp.* Root329 is the major isolate responsive to lack of coumarin biosynthesis among members of the retrievable microbial community.

**Figure S8.**
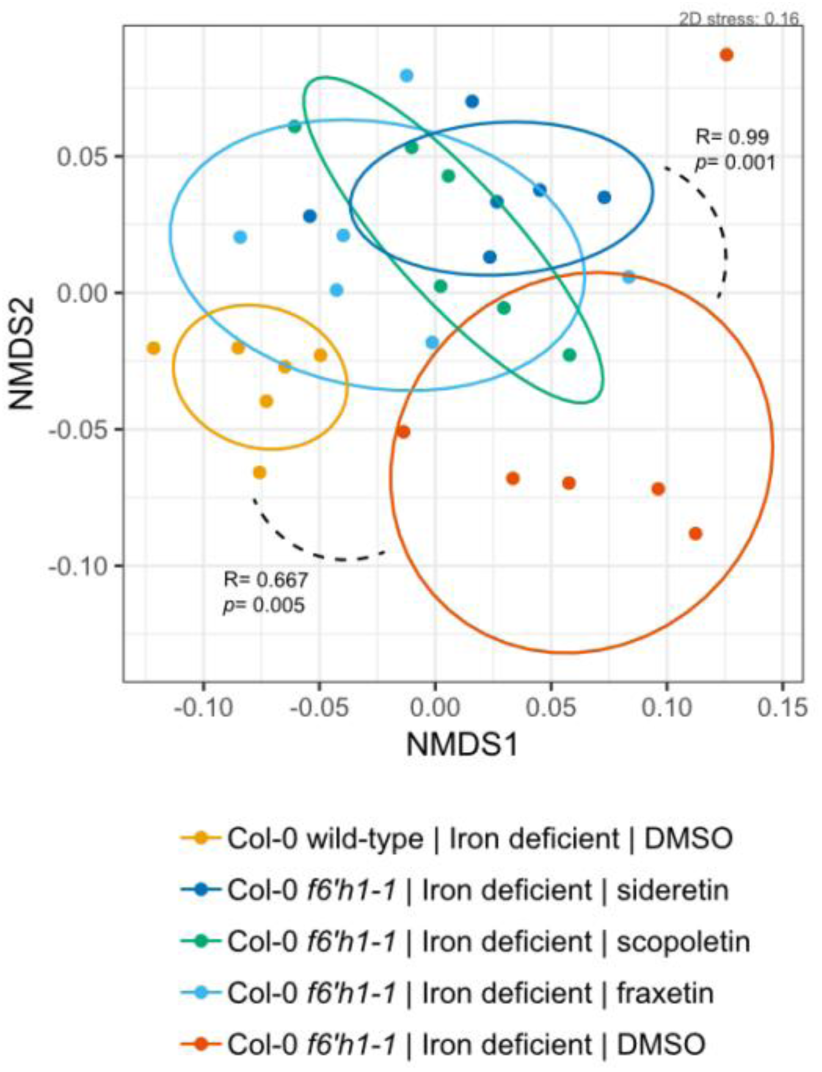
Shifts in root community profiles of wild-type phenocopy experiments in *f6’h1*. Non-metric multidimensional scaling (NMDS) on Bray-Curtis dissimilarity matrices of microbial community profiles at the wild-type, *f6’h1-1,* and *f6’h1-1* coumarin-added roots in iron-deficient media (of 20-day-old seedlings, incubated with the SynCom for 7 days). Addition of oxidized coumarins noticeably shifts the microbial community profile, possibly towards that of wild-type roots (*f6’h1* plants present with wild-type phenocopy). All other descriptors are the same as in Figure S3.

**Figure S9.**
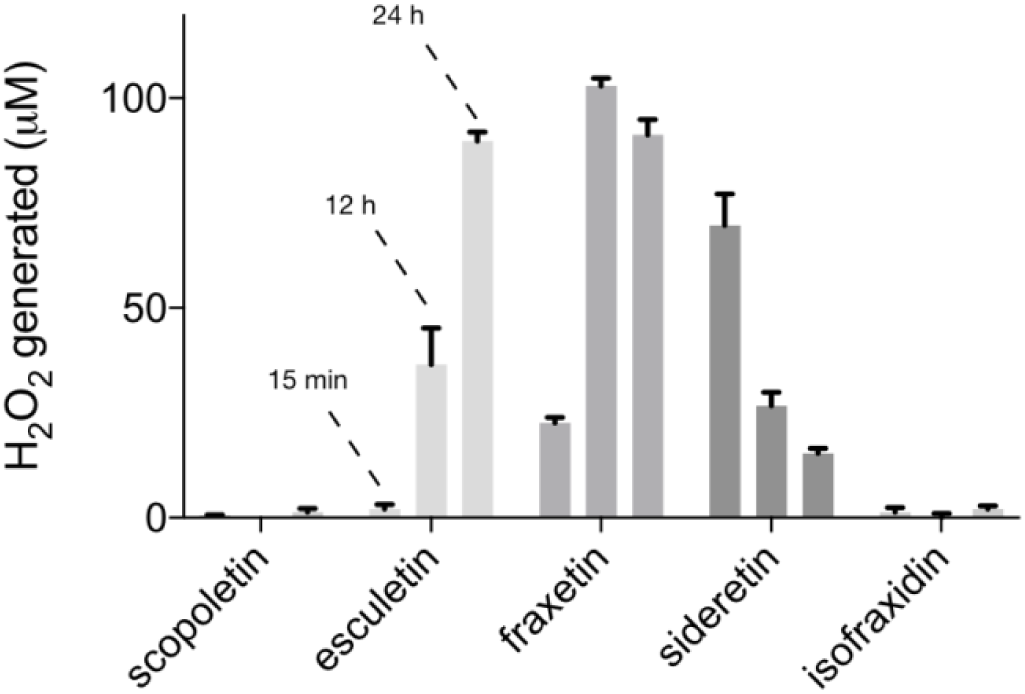
Catecholic coumarins generate hydrogen peroxide in plant growth media. Bar graph depicting a 24-hour time-course of hydrogen peroxide generated by 100 µM of each coumarin indicated in iron-deficient Murashige and Skoog plant growth media. Hydrogen peroxide levels was assayed by the Ferrous Oxidation-Xylenol orange (FOX) method. The mean and standard deviation of 4 replicates is shown.

**Figure S10.**
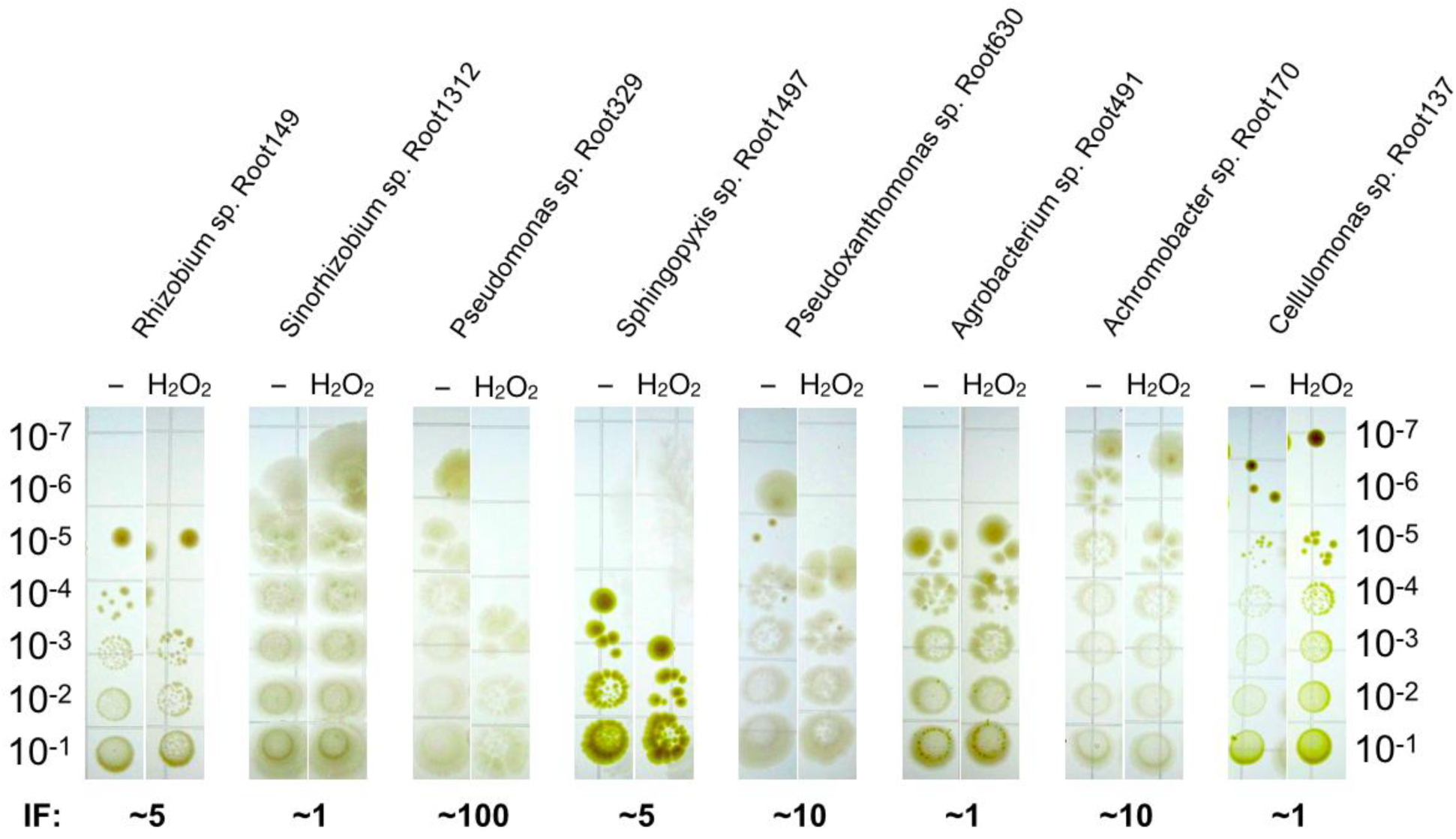
Survival after peroxide challenge of SynCom members. SynCom strains at the same density (OD_600_ 0.5) were stressed with 5 mM hydrogen peroxide for 1 hour. Survival after the hydrogen peroxide challenge was determined by CFU counts. The inhibition factor (IF) was estimated as the factor with which peroxide inhibited survival of the strain compared to the control (–). *Pseudomonas sp.* Root329 is among the SynCom members most sensitive to hydrogen peroxide.

**Figure S11.**
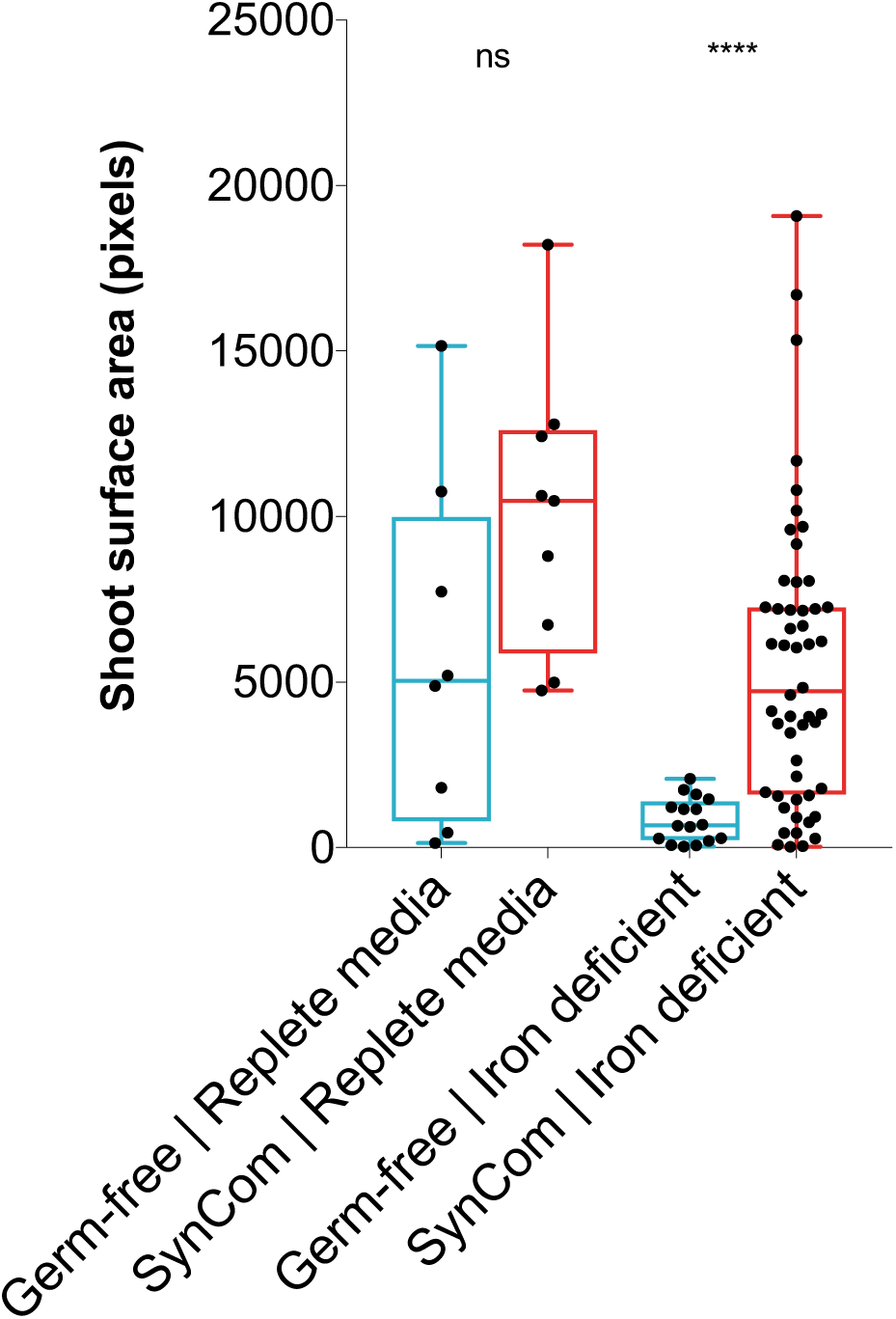
*A. thaliana* seedling growth promotion by the SynCom. Wild-type seedling growth promotion by the SynCom. Each data point represents the shoot surface area of a seedling in pixels (as determined by image processing using imageJ script). Number of planted seeds per condition. In this assay condition, buffering at circumneutral pH causes the Fe(III)-EDTA chelate to precipitate out of solution, requiring the mobilization of Fe^3+^ for seedling growth and development. Half-strength Murashige and Skoog media buffered to pH 7.1 by 10 mM HEPES was used, with addition of 10 µM Fe-EDTA (iron deficient), or 100 µM Fe-EDTA (replete). Significance testing by Mann-Whitney using PRISM (GraphPad).

**Figure S12.**
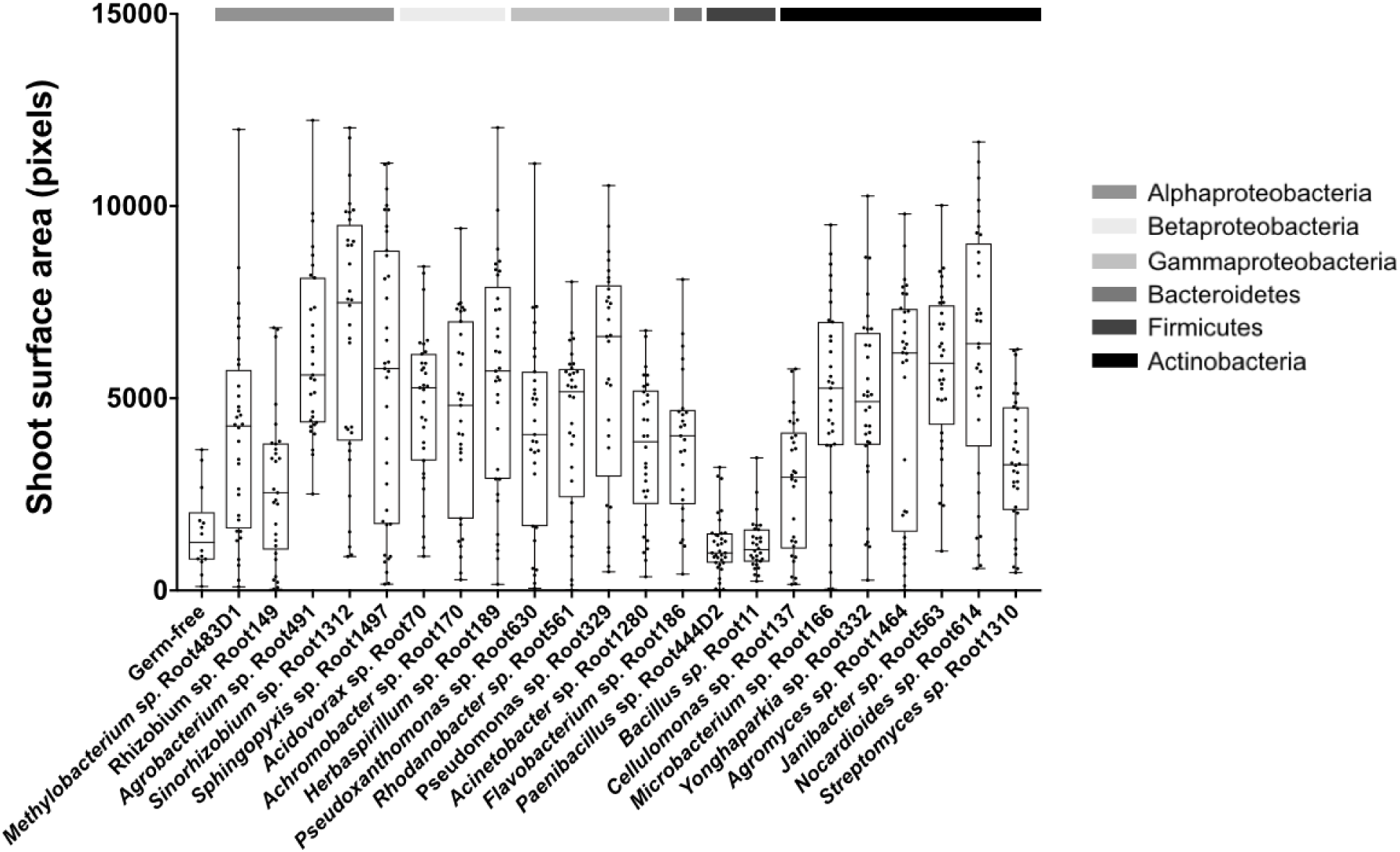
*A. thaliana* seedling growth promotion by SynCom members. Each data point represents the shoot surface area of a single seedling (that germinated) on iron-deficient half-strength Murashige and Skoog media (20 µM Fe-EDTA, 10 mM HEPES-buffered to pH 7.1). Box plots with median and interquartile ranges are shown. All isolates could be cultured from the root after the experiment, except for the Firmicutes. The number of planted seeds per condition is 32.

**Figure S13.**
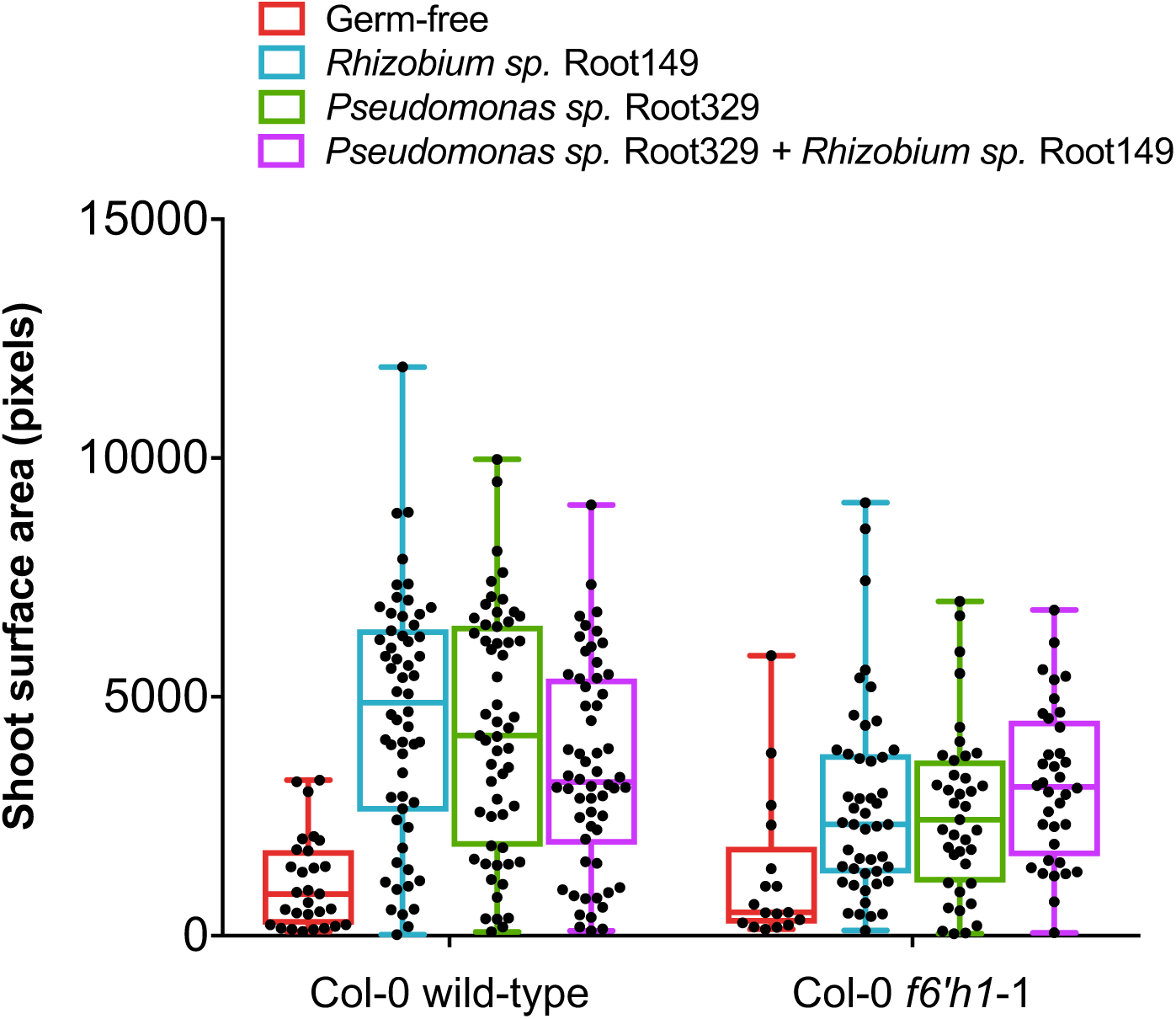
Wild-type and *f6’h1* seedling growth promotion by SynCom members under iron-deficient conditions. Each data point represents the shoot surface area of a single seedling (that germinated). Box plots with median and interquartile ranges are shown. For each condition, 64 seeds were planted onto half-strength HEPES-buffered MS growth media buffered to pH 7.1 containing the iron concentration indicated and inoculated with or without the indicated isolate(s). The catecholic coumarin-sensitive *Pseudomonas sp.* Root329 promoted the growth of wild-type and *f6’h1* seedlings, equally.

## Materials and Methods

### SynCom strains and plant lines

Bacteria and plant lines used in this study are listed in Tables S1 and S2. To design the SynCom used in this study we first extracted (*in silico*) the V5-V7 variable regions from the 16S rRNA of the 206 At-RSPHERE genome-sequenced root isolates^21^. We next developed a custom script in MATLAB (MathWorks) that randomly picked strains for which V5-V7 regions were at most 97% similar to one-another, while maximizing for the number of phylogenetic families represented in the SynCom. This resulted in a SynCom composed of 27 strains that was reduced to 22 strains after (1) confirming with Sanger sequencing that the V5-V7 16S rRNA region from each strain fully matched with that of the sequenced genome, and (2) selecting for strains that reached saturation in quarter-strength TSB within 5 days (for experimental considerations).

### Plant growth conditions

*A. thaliana* seeds were surface sterilized for 5 minutes in a 70% ethanol solution, followed by 15 minutes in 50% bleach solution, washed three times with sterile water, and stratified at 4°C for 3 days before planting. Circular disks 3 cm in diameter were cut out from a 0.025 x 0.005 opening size PTFE mesh (McMaster-Carr) and sterilized by autoclaving. 3 mL of 1x Murashige and Skoog (MS) medium (100 µM Fe-EDTA, 10 mM HEPES, pH = 7.1; iron sufficient condition) was added to the wells of a 6-well plate (Corning), a PTFE mesh raft was placed on the medium surface in each well, and 16 seeds were individually planted in a square grid on the surface of the raft with a pipette. The plates were wrapped in aluminum foil to etiolate (thereby reduce wetting of the shoots) for 3 days. The plants were grown in a growth chamber under a 16 h light cycle (photon flux of 100 µmol/m^2^/s, 22°C, 50% relative humidity). 13 days after planting, the seedling roots were washed with 1x phosphate-buffer saline (PBS) and the medium was replaced with 3 mL of either iron sufficient (regular) 1xMS (100 µM Fe-EDTA) or iron-deficient (no added iron) 1xMS inoculated with the SynCom.

### Root inoculation and harvest

Bacterial strains were cultured to saturation (RT, 14 mL round-bottom Falcon tubes, shaken at 250 rpm) in 4 mL of quarter-strength tryptic soy broth (TSB) for 5 days, spun down, and washed 3 times with 1xPBS. OD_600_ of each strain solution was determined using a BioTek plate reader, and the final OD_600_ of all strains was normalized by dilution with 1x PBS in the final mixture containing the 22-member SynCom. 1xMS media (with or without added iron, or coumarins) was inoculated to a final composite OD_600_ of 0.005–0.01. Following inoculation, plants were grown for an additional 7-8 days under a 16 h light cycle at 22°C until harvest. Plant roots were harvested by washing roots in 10 mL 1xPBS, followed by excision of the root from the shoot approximately 5 mm below the rosette, and storage in lysing matrix E tubes (MP Biomedicals) at −80°C prior to total DNA extraction. For the purposes of downstream processing and data representation we considered 2 wells (containing total 32 planted seeds) a technical replicate. A biological replicate is defined as an experimental batch with independently prepared seed and SynCom members re-inoculated from 40% glycerol stocks stored at −80°C.

### DNA extraction and CFU counts

All frozen samples were homogenized twice for 30 seconds at 5.0 m/s using a Precellys24 tissue lyser (Bertin Technologies). DNA from root tissue and inocula were harvested separately using the FastDNA SPIN Kit for Soil (MP Biomedicals). From this step forward, all DNA concentrations were determined using the PicoGreen dsDNA Assay Kit (Life Technologies).

For colony-forming unit (CFU) counts in plant growth media, 200 µL of media was 10-fold serially diluted 7 times in 1x PBS, followed by spotting of 5 µL of each serially-diluted sample onto quarter-strength TSB-agar. Plates were left at RT for 5 days to allow for growth sufficient of all strains before imaging.

### 16S rRNA community profiling

16S rRNA amplicon generation and library preparation were performed according to Bai et al. (2015)^21^. Sample DNA was diluted to 3.5 ng/µL prior to 16S rRNA amplicon PCR by degenerate primers 799F and 1193R. These primers target the V5-V7 variable region of the 16S rRNA locus. Each sample was amplified in triplicate. The 25 µL PCR mix contained 1x incomplete buffer (Invitrogen), 2 mM MgCl_2_ (Invitrogen), 0.3% BSA (New England Biolabs), 200 µM dNTPs (Invitrogen), 300 nM of each primer, 2 U of Taq DNA polymerase (Invitrogen) and 10 ng of template DNA. The PCR program was as follows: a denaturation step at 94°C for 2min, and 25 cycles of 94°C for 30s, 55°C for 30s and 72°C for 60s, followed by a final elongation step of 5 min at 72°C. The triplicate PCR reactions were pooled and treated with 20 U exonuclease I and 5 U Antarctic phosphatase (New England Biolabs) for 30 mins at 37°C. Following heat-inactivation at 85°C, the digested mixture was used as template for the barcoding PCR reaction. We used the Illumina-compatible primers B5-F and 1 of 96 differentially barcoded reverse primers (B5-1 to B5-96, Supplementary Data).

Following 10 cycles of PCR (in triplicate from the pooled digests) using the same program as described above, the triplicates were pooled, and run on a 1% agarose gel to separate bacterial amplicons from root mitochondrial amplicons. All samples were extracted using the QIAquick Gel Extraction Kit (Qiagen) according to the manufacturer’s instructions. Following DNA concentration measurements by PicoGreen dsDNA Assay Kit (Life Technologies), 100 ng of each barcoded sample were pooled. The pooled amplicon library was purified using the Agencourt AMPure XP Kit (Beckman Coulter) and subjected to sequencing on the Illumina MiSeq platform (Illumina Inc.) using the MiSeq Reagent kit v3 (2×300 bp paired-end).

### Processing and statistical analysis of 16S rRNA counts

The Quantitative Insights into Microbial Ecology (QIIME) pipeline was used to process raw sequencing data^37^. The data was demultiplexed and mapped to known 16S rRNA sequences of the 22-member SynCom using the *“pick_closed_reference_otus*” function (see Supplementary Data for reference sequences used). We included *A. thaliana* mitochondrial and plastid DNA sequences to the alignment to account for contamination. To account for errors in sample preparation, sequencing, or contamination, samples with an unusually high abundance of unassigned reads (more than 2.5-fold the interquartile range from the median) were omitted from downstream processing. BIOM tables obtained from QIIME were used as input in the R statistical environment for statistical analysis and plotting (R Development Core Team; http://www.R-project.org). Ordination analysis by non-metric multidimensional scaling (NMDS) was primarily used to visualize Bray-Curtis dissimilarity of the community profiles between samples after rarefying the data (to 10^4^ reads per sample, accounting for differences in sequencing depth between samples)^38^. Statistical significance was determined using Analysis of Similarity (ANOSIM), i.e. a non-parametric rank-ordered statistical test with a significance threshold set to *p* = 0.01. The level of dissimilarity between groups (R values) are scaled to between −1 and +1, where 1 indicates samples between groups are more dissimilar than within groups, 0 indicates no relationship between the groups, and −1 indicates a higher degree of similarity between the groups than within the groups (here, groups are defined as replicates of a single condition or plant genotype).

In Figures 2b, 3b and 4a, Z scores were calculated from read counts obtained after rarefaction to 10^4^ for every sample. The differences between read counts for isolates from the roots of mutant lines in each sample (*x*i_mutant) grown wild-type roots (μ_i_WT_) was normalized to the standard deviation (σ_i_WT_) of read counts for and corresponding isolates from synchronously-wild-type root isolates (Z = *x*_i_mutant_ μ_i_WT_)/ σ_i_WT_). Variable colonization of the root was observed for *Sphingopyxis sp.* Root1497, as it was always present in the 16S rRNA reads of the inoculum, but not consistently in reads of the root between independent experiments, such as in Fig. 4a. Z scores for strains were only calculated in experiments for which all strains were consistently detected above the read threshold after rarefaction (≥2).

### Coumarin-mediated growth inhibition assays

Growth-inhibition by coumarins was determined by inoculating 25 mL molten LB-agar (aat 40°C) to a final OD_600_ 0.05 with the bacterial strain of interest (washed with 1x PBS), followed by placing a 0.5 x 0.5 cm sterile filter disc impregnated with 5-10 mM of the compounds of interest on top of the solid agar. The degree of growth inhibition was determined visually by the radius of growth inhibition (dark halo surrounding the disc) after 48 h at 30°C.

### Ferrous oxidation-xylenol orange (FOX) assay for peroxide generation by coumarins

100 µM final concentration of each coumarin – scopoletin, esculetin, fraxetin, sideretin or isofraxidin – was added to 10 mM HEPES-buffered half-strength MS media at pH 7.1 without iron added. Hydrogen peroxide generation was monitored by FOX reagent over a 24-hour period at 560 nm absorbance using a BioTek plate reader (as compared to a standard curve generated using calibrant hydrogen peroxide dilution). In the FOX assay, hydrogen peroxide in the sample reacts with ferrous iron to form ferric iron which complexes with xylenol orange to form a light-orange color. Peroxide generation by tested coumarins was confirmed orthogonally using peroxide test sticks (Quantofix).

### Peroxide challenge of SynCom strains

SynCom isolates were cultured to saturation in quarter-strength TSB (5 days), washed with 1x PBS, normalized to an OD_600_ of 0.25 in 10 mM HEPES-buffered quarter-strength TSB, and subsequently subjected to 5 mM freshly-prepared hydrogen peroxide (with H_2_O as control) for 1 hour at RT without shaking, before dilution-spotting onto quarter-strength TSB solid media. Images of colonies were taken after 5 days of incubation at RT.

### Promotion of A. thaliana seedling growth by SynCom members under iron-deficient conditions

We designed an assay to quantitative assess the degree of bacteria-mediated growth promotion of *A. thaliana* seedlings. In this assay condition, buffering at circumneutral pH causes the Fe(III)-EDTA chelate to precipitate out of solution, requiring the mobilization of Fe^3+^ for seedling growth and development. Each strain was inoculated at an OD_600_ of 0.005 into 10 mM HEPES-buffered half-strength MS media at pH 7.1 containing either 20 µM Fe-EDTA (iron-deficient condition for wild-type), 50 µM Fe-EDTA (iron-deficient condition for *f6’h1*), or 100 µM Fe-EDTA (replete media condition for wild-type and *f6’h1*). 16 seeds were planted onto each plate containing 30 mL of MS-agar using an evenly-spaced 4×4 grid. Seeds were subsequently etiolated for 3 days and grown at 22°C with a 16/8-hour light/dark cycle (photon flux of 100 µmol/m^2^/s, 50% relative humidity) for 14 days prior to imaging. The degree of growth promotion was quantified as shoot surface area, which was assessed by demarcating shoots in the image, and subsequently determining the number of pixels corresponding to shoots using a custom ImageJ (NIH) script.

